# Critical neuronal avalanches arise from excitation-inhibition balanced spontaneous activity

**DOI:** 10.1101/2025.11.17.688775

**Authors:** Maxime Janbon, Mateo Amortegui, Enrique Carlos Arnoldo Hansen, Sarah Nourin, Virginie Candat, Germán Sumbre, Adrián Ponce-Alvarez

## Abstract

Neuronal avalanches are sequences of neural activations exhibiting scale-invariant statistics, indicative of critical dynamics. Theoretical studies proposed that the balance between excitation (E) and inhibition (I), along with neuromodulation, are key factors influencing this critical behavior. Here, we performed in-vivo studies to investigate the role of E and I neurons in generating neuronal avalanches in the optic tectum of zebrafish larvae. For this, we used double-transgenic zebrafish larvae expressing cell-type-specific fluorescent proteins and GCaMP6f, combined with immunostaining and selective-plane illumination microscopy to monitor spontaneous neuronal activity and neurotransmitter identity. We found that neural activity approached criticality at balanced and slightly excitatory-dominated E-I ratios but became disordered when E-I ratios were imbalanced. A stochastic network model operating at a critical point, where excitation and inhibition couplings are balanced and balanced amplification drives network avalanches, successfully reproduced the observed statistics of neuronal avalanches and their dependence on E-I ratio fluctuations.

## INTRODUCTION

Neural systems can display cascading collective activity patterns, known as neuronal avalanches, characterized by sequences of neuronal activations. Neuronal avalanches have been observed at multiple spatiotemporal scales using a variety of recording techniques, including multi-electrode arrays^1–6^, optical imaging^7^, and even low– temporal-resolution recordings such as fMRI^8,9^, and calcium imaging^10–12^. The distributions of sizes and durations of neuronal avalanches indicate scale-invariance, i.e., they follow power-law statistics with precise power exponents, a shared feature with other critical systems producing cascading events, e.g. earthquakes, rock fractures, and many others^13^. The observation of power-law statistics in neural activity has contributed to support the idea that spontaneous neural activity operates close to a phase transition^14^. Moreover, at critical points, computations of neural systems are optimized in terms of information transmission, storage, processing, efficient coding, energy efficiency, and the response sensitivity-reliability tradeoff^11,15–22^.

Critical dynamics are thought to arise through self-organization, enabling the system to adjust its parameters and converge automatically to the critical point. It has been suggested that neural systems achieve self-organization via simple homeostatic plasticity rules that rewire the neural network, effectively tuning the system’s parameters toward their critical values^23–28^. The balance between excitation and inhibition (E-I balance), along with neuromodulation, is commonly proposed as key control parameter that is changed during the self-organization process^21,29–31^. The importance of E-I balance in shaping the statistics of neuronal avalanches—i.e., their size, duration, and occurrence times— has been suggested by theoretical models^32–35^ and in vitro pharmacological experiments^36^. However, how the dynamics of excitatory (E) and inhibitory (I) neurons evolve and interact during neuronal avalanches and scaling behavior remains unknown. Here, we studied the dynamics of excitatory and inhibitory neurons during spontaneous avalanche activity in the zebrafish optic tectum.

Previous studies using immunostaining and in situ hybridization have indicated that most tectal neurons are glutamatergic or GABAergic, with few cholinergic neurons^37–40^ and the absence of aminergic (noradrenergic and serotonergic) and glycinergic neurons at this developmental stage^41,42^. In the present work, we detected these cell-types and studied the contribution of the spontaneous activity of E and I neurons to neuronal avalanches in the optic tectum of the zebrafish. To study the activity of these cell types, we used double-transgenic zebrafish larvae expressing the genetically encoded calcium indicators GCaMP6f and mCherry under the Vglut2a promoter (glutamatergic), in combination with immunostaining to label GABAergic (gad1b) and cholinergic (ChAT) neurons and selective-plane illumination microscopy (SPIM) to monitor the neuronal dynamics. Finally, a network model composed of E and I stochastic neurons was built to understand the underlying mechanisms.

Our results show that spontaneous fluctuations in excitatory and inhibitory activity influenced neuronal avalanche statistics in the zebrafish optic tectum. Neuronal avalanches approached criticality when excitatory and inhibitory activity were balanced but deviated when E-I ratios were imbalanced. Notably, the model accurately captured the observed avalanche statistics and their sensitivity to E-I fluctuations around a critical point defined by excitatory and inhibitory synaptic strengths.

## RESULTS

### Spontaneous activity of the optic tectum and neuronal avalanches

To study the role of the different cell types in the generation of the neuronal avalanches, we analyzed spontaneous neuronal activity from the optic tectum of 10 zebrafish larvae (5-8 days post-fertilization, dpf) using SPIM (see STAR **Methods**). We used double-transgenic zebrafish in combination with immunostaining to identify glutaminergic (excitatory, E), GABAergic (inhibitory, I), and cholinergic (Ch) neurons — the three types of neurons found in the optic tectum^37–42^ (**Figure 1A**). Spontaneous neural activity was recorded during 1h at 15Hz (therefore the time bin corresponds 66.67 ms). The total number of recorded neurons per fish was 2,000 and the average proportion of neurons from the identified cell types E, I, and Ch, was 55.2%, 43.6%, and 1.2%, respectively (see **Table S1**). To detect calcium events, the fluorescence signal of each neuron was binarized by imposing a threshold (see STAR **Methods**). The average activation rate was 0.18 ± 0.02 Hz, 0.16 ± 0.02 Hz, and 0.18 ± 0.02 Hz, for cell types E, I, and Ch, respectively. Given the small number of recorded Ch neurons, the remainder of this study focuses on E and I activity.

**Figure 1.**
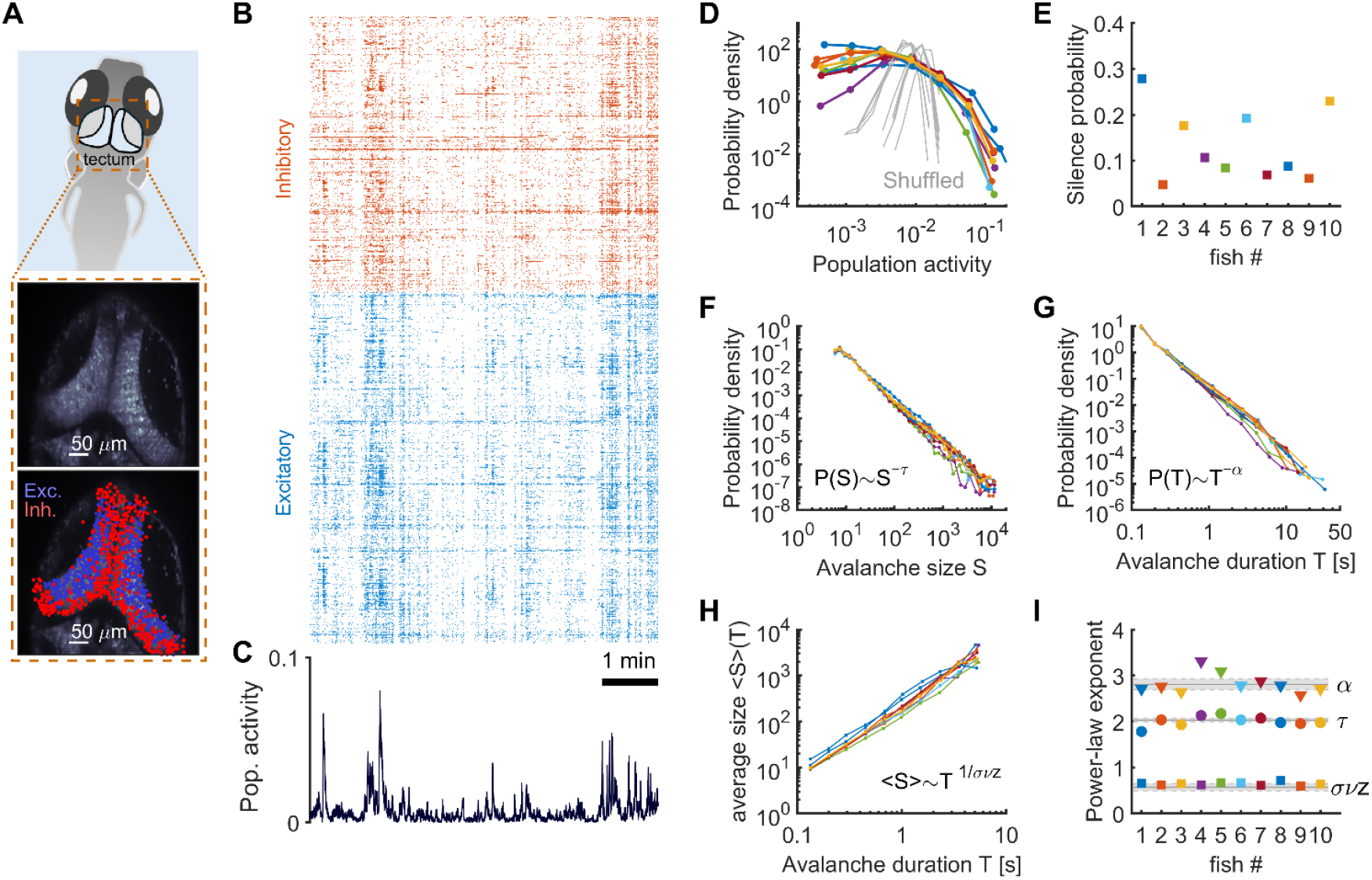
Spontaneous optic tectum activity. **A)** Double-transgenic zebrafish in combination with immunostaining to identify glutaminergic (excitatory, E) and GABAergic (inhibitory, I) neurons in the optic tectum. The average image of a recorded SPIM plane is shown (*middle*) together with the recorded neurons belonging to different cell types (*bottom*). **B)** Sample of collective binary activity, for an example fish. The activations of neurons identified as excitatory or inhibitory cells are shown in *blue*, or *red*, respectively. **C)** Population activity, i.e., proportion of active neurons. **D)** Probability density of the population activity, for the different fish (colored traces). The gray distributions correspond to the probability density of the population activity when the data is shuffled in time. **E)** Probability of observing a time bin with zero population activity, i.e., no activation from any neuron, for each fish. **F-H)** The probability distributions of avalanche sizes (*S*), avalanche durations (*T*), and the relation between ⟨*S*⟩ and *T*, were fitted by power laws *P*(*S*)∼*S*^−*τ*^, *P*(*T*)∼*T*^−*α*^, and ⟨*S*⟩(*T*)∼*T*^1⁄*σνz*^. **I)** The measured power-law exponents for each fish are shown. Triangles: *α* exponent; circles: *τ* exponent; squares: *σνz* exponent. Estimation errors are smaller than the size of the symbols. The gray horizontal lines and the gray shaded areas indicate the expected critical exponents and their uncertainty, respectively, in random field Ising models. Individual power-law exponents, estimation errors, and power-law tests are detailed in **Table S1**.

Spontaneous neuronal activity exhibited large, correlated events where multiple neurons activated simultaneously (**Figure 1B**). Population activity showed pronounced fluctuations and extended silent periods (**Figure 1C-E**), reflecting transient correlated dynamics among the neurons, indicative of neuronal avalanches.

Neuronal avalanches consist of sequential co-activations of nearby neurons, characterized by their duration *T* and size *S* (number of neuronal activations) (see STAR **Methods**; note that the definition of avalanches imposes a spatial constraint). The size and duration distributions followed approximate power-law behavior, i.e., *P*(*S*)∼*S*^−*τ*^ and *P*(*T*)∼*T*^−*α*^ (**Figure 1F-G**), with average power-law exponents equal to ⟨*τ*⟩ = 2.01 ± 0.03 and ⟨*α*⟩ = 2.82 ± 0.07. Moreover, the average size ⟨*S*⟩(*T*) of avalanches of duration *T* was given by the scaling relation ⟨*S*⟩(*T*)∼*T*^1⁄*σνz*^ (**Figure 1H**), with an average power-law exponent equal to ⟨*σνz*⟩ = 0.64 ± 0.01. The measured exponents {*α, τ, σνz*} closely matched predictions for systems displaying critical avalanche dynamics (*τ* = 2.03 ± 0.03 and *α* = 2.81 ± 0.11), derived from numerical simulations of the random field Ising model (RFIM) — the paradigmatic model of disorder-induced phase transitions^13,43^ that was used to compared exponents in previous work^11^ (**Figure 1I**; see **Table S1** for detailed values, uncertainties, and power-law tests). Average relative deviations of the exponents with respect to their mean and to the values predicted by the model are 4.6% and 7.6%, respectively, indicating a high degree of robustness. Furthermore, it is known that the critical exponents *α, τ*, and *σνz* must obey the relation^5,13,43^ (*τ* − 1)⁄(*α* − 1) = *σνz*, called crackling-noise relation. We evaluated the deviation from this relation using the following index^44^: DCC = |*σνz*^pred^ − *σνz*|, where *σνz*^pred^ = (*τ* − 1)⁄(*α* − 1), and found that the average deviation was small and equal to 0.09. Neuronal avalanche statistics differed from those of shuffled datasets and remained robust over a wide range of avalanche detection and fitting parameters (see STAR Methods; see **Supplemental Figures S1**−**2**).

To facilitate comparison with previous studies, we also computed neuronal avalanches without spatial constraints, defining them as periods during which the summed activity exceeded a threshold (see STAR **Methods**), following approaches used in earlier work^1–6,45,46^. This yielded power-law exponents equal to ⟨*τ*⟩ = 1.61 ± 0.04, ⟨*α*⟩ = 1.64 ± 0.02, and ⟨*σνz*⟩ = 0.87 ± 0.03 (DCC = 0.08; see **Supplemental Figure S3**, and **Table S2** for details). These values were consistent with previous studies using the spatially unconstrained definition of neuronal avalanches and indicative of criticality^47–50^.

Furthermore, we studied the contribution of excitatory and inhibitory activity in avalanche activity. As a preliminary step, we examined whether stationary pairwise correlations were cell-type specific and could influence avalanche dynamics. Yet, this analysis revealed neither cell-type-specific modularity nor differences in network centrality between excitatory and inhibitory neurons (see **Supplemental Note S1** and **Supplemental Figure S4**). These results suggest that neuronal avalanches are temporally transient, non-stationary events that may involve higher-order correlations beyond static (and zero-lag) pairwise interactions.

### Statistics of neuronal avalanches as a function of E-I ratio

To study the influence of excitation and inhibition on the avalanche dynamics in the optic tectum, we examined the relationship between the statistics of neuronal avalanches and the relative spontaneous activity of excitatory and inhibitory neurons by calculating the global excitation-inhibition (E-I) ratio of the observed network of neurons. For each recording session, the E-I ratio was defined as: *r*_*E*_ (*t*)⁄[*r*_*E*_ (*t*) + *r*_*I*_(*t*)], where *r*_*E*_ (*t*) and *r*_*I*_(*t*) represent the activity (i.e., proportion of active neurons) of tectal excitatory and inhibitory neurons at time *t*, respectively (**Figure 2A-B**). We observed that the E-I ratio displayed temporal fluctuations around the mean value of 0.51 ± 0.18 (std.) (**Figure 2C**), with relatively strong correlations between *r*_*E*_ (*t*) and *r*_*I*_(*t*) (**Figure 2D**), and that the autocorrelation of such fluctuations decayed after a few seconds (**Figure 2E**), a timescale that was relatively longer than the durations of neuronal avalanches. Indeed, the average duration of avalanches was 284 ms and 99% of the neuronal avalanches were shorter than 2.4 s. In the following we studied the dependence of neuronal avalanches on the E-I ratio during which they occurred.

**Figure 2.**
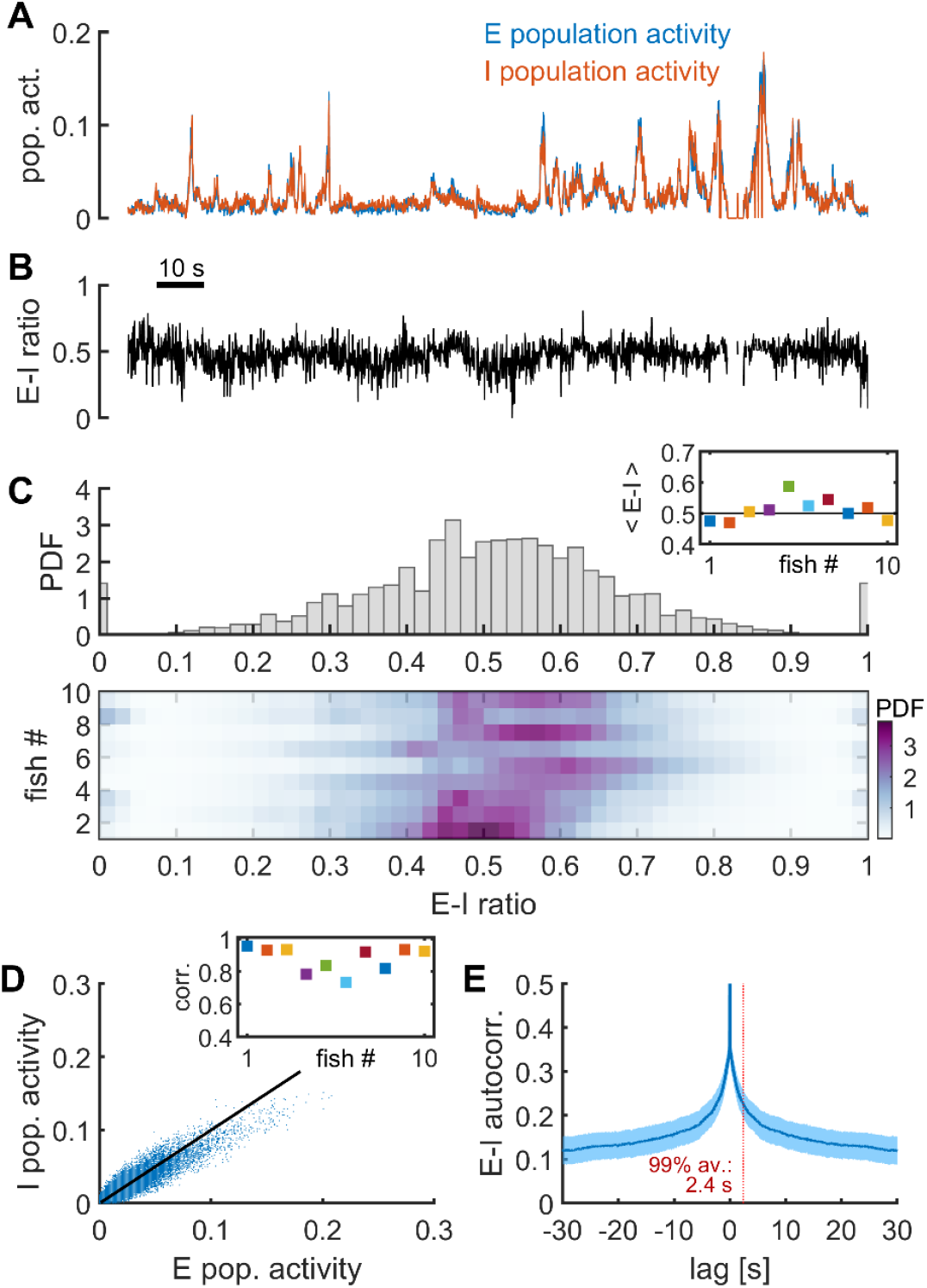
Excitation-inhibition ratio. **A)** Excitatory and inhibitory population activities, for an example fish. **B)** Fluctuations of the E-I ratio of the example fish in (A). **C)** Probability density function of the E-I ratio across all fish (*top*) and for each fish separately (*bottom*, colormap). **D)** Correlation between excitatory and inhibitory population activities for one example fish. *Inset:* correlation coefficients for the different fish. **E)** The average autocorrelation of the E-I ratio. Note that the decay of the autocorrelation is longer than the duration of most (99%) of the neuronal avalanches, which are shorter than 2.4s. The shaded area indicates SEM.

We observed that the size and the number of neuronal avalanches tended to decrease during periods in which E-I deviated from its mean value (**Figure 3A**). To quantify this, we associated each detected neuronal avalanche with the average E-I ratio between its initiation (at time *t*_1_) and termination (at time *t*_2_). For each fish, we grouped the E-I ratios in three categories: low E-I ratios (lowest 25%), balanced E-I ratios (E-I ratios within the 25^th^-75^th^ percentiles), and high E-I ratios (highest 25%). We found that the neuronal avalanches were larger, longer and more diverse when the E-I ratio during the avalanche was close to balanced values (**Figure 3B-C**). This was shown by a finer partitioning of the range of E-I ratios into bins representing the deciles of the E-I ratio distribution (i.e., each bin contained one-tenth of the sample, to avoid sampling biases). Using this binning, we calculated the average size ⟨*S*⟩ and average duration ⟨*T*⟩ of the neuronal avalanches as a function of the E-I ratio. We found that both ⟨*S*⟩ and ⟨*T*⟩ quantities peaked around the balanced and slightly excitatory-dominated E-I ratio (**Figure 3D-E**).

**Figure 3.**
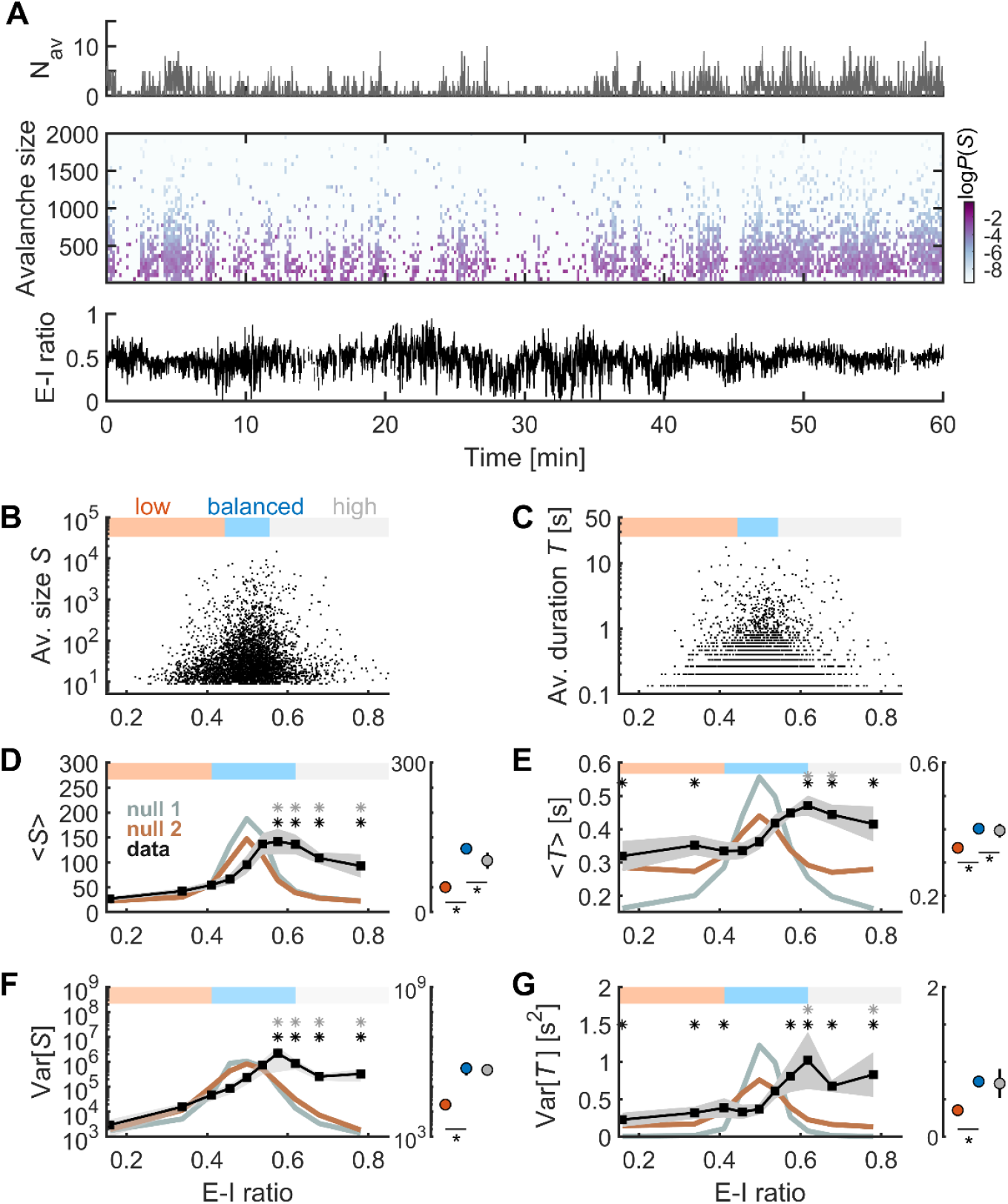
The statistics of neuronal avalanches as a function of the E-I ratio. **A)** *Top:* Number of neuronal avalanches per time bin. *Middle:* Probability density of the size of neuronal avalanches observed in each time bin. *Bottom:* Fluctuations of the E-I ratio. **B-C)** Size (B) and duration (C) of neuronal avalanches as a function of the average E-I ratio during the avalanche, for an example fish. Each dot represents a neuronal avalanche. The E-I ratios were grouped in the categories: low (*red*), balanced (*blue*), and high (*gray*), delimited by the 25th-75th percentiles of the distribution of E-I ratios for each fish. **D-G)** *Left panels:* Average size (D) and duration (E), as well as they respective variances (F-G), of neuronal avalanches for different bins of E-I ratios (i.e., deciles of the E-I ratio distribution from all fish). Gray patches indicate SEM. Gray and golden traces indicate the expected values of null models 1 and 2, respectively. Black asterisks: significantly higher values with respect to null model 1 (shuffling E-I ratio time-courses, *p* < 0.05). Gray asterisks: significantly higher values with respect to null model 2 (shuffling neuronal identities, *p* < 0.05). *Right panels:* Avalanche size, duration, and their respective variances for avalanches occurring at low, balanced, and high E-I ratios. *: *p* < 0.05, post hoc t tests following one-way Kruskal-Wallis ANOVA.

Given that, on average, large-size neuronal avalanches were longer (since ⟨*S*⟩(*T*)∼*T*^1⁄*σνz*^), it could be that the large ⟨*S*⟩ and long ⟨*T*⟩ around the balanced E-I ratio resulted from the estimation error of the mean E-I ratio, i.e., the E-I ratio approached its mean value over long periods (i.e., *t*_2_ − *t*_1_). To rule this possibility out, we compared the values of ⟨*S*⟩ and ⟨*T*⟩ with those obtained from a data-driven null model in which the time series of the E-I ratio was shuffled in time (2,000 repetitions). This surrogate dataset (null-model 1) preserved the averaging effect but decoupled the timing of neuronal avalanches from the E-I ratio. We found that, for some balanced and all excitatory-dominated E-I ratios, neuronal avalanches were significantly larger and longer than those in the null model (*p* < 0.05, permutation test; **Figure 3D-E**, black asterisks). Additionally, we further tested the significance of the relationship between neuronal avalanches and the E-I ratio using a second control dataset (null-model 2), where the identities of the neurons were randomly permuted (2,000 permutations). This second null model preserved the statistics of the neuronal avalanches and overall network activity but removed information regarding neuron cell types. In this second control, we found that ⟨*S*⟩ and ⟨*T*⟩ were significantly larger than obtained after randomizing the cell identities for some of the balanced and excitatory-dominated E-I ratios (*p* < 0.05, permutation test; **Figure 3D-E**, gray asterisks).

Similarly, we found that the variance of size and duration of neuronal avalanches, *var*[*S*] and *var*[*T*], were maximal around balanced and slightly excitatory-dominated E-I ratios, and significantly larger than those obtained from the two null models controlling for the averaging effect and randomized cell identities, respectively (*p* < 0.05, permutation test; **Figure 3F-G**). Altogether, these results indicate that neuronal avalanches were maximally larger, longer and more diverse in periods when the network’s excitation and inhibition balanced each other or were slightly dominated by excitation.

We then analyzed the distributions of the sizes and durations of neuronal avalanches for each E-I ratio category. We found that the distributions could be approximated by power laws for all E-I ratio categories, but the distributions for avalanches occurring at low or high E-I ratios decayed faster than those at balanced E-I ratios (**Figure 4A-D**). Consequently, the effective power-law exponents describing the distributions of sizes and durations of neuronal avalanches were significantly higher for low and high E-I ratios with respect to the balanced E-I ratios (for exponent *τ*: *F*_(2,27)_ = 7.35, *p* = 0.003; for exponent *α*: *F*_(2,27)_ = 7.94, *p* = 0.002; one-way ANOVA, **Figure 4E**). Furthermore, we calculated the avalanche distributions for different bins of E-I ratios, that were chosen to contain an equal number of neuronal avalanches (1,000), and estimated the power-law exponents *τ, α*, and *σνz* for each bin (**Figure 4F-H**). We found that the average power-law exponents approached the model’s critical values for balanced E-I ratios, while they increased for imbalanced E-I ratios.

**Figure 4.**
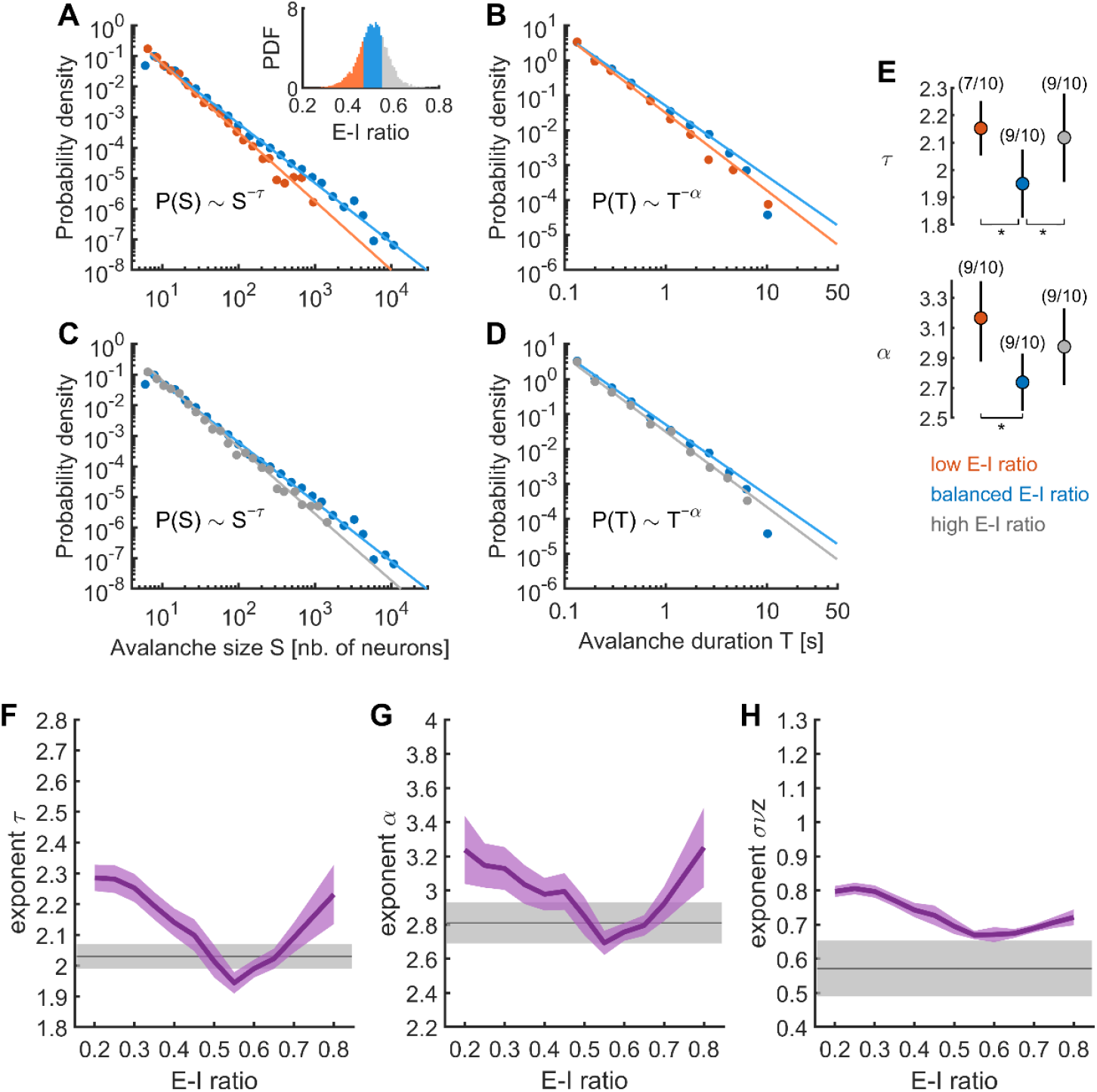
Neuronal avalanche exponents as a function of the E-I ratio. **A)** Distribution of avalanche sizes (*S*) for avalanches occurring at low E-I ratios (*red*, lowest 25% E-I ratios) and at balanced E-I ratios (*blue*, E-I ratios within the 25^th^-75^th^ percentiles). Solid lines indicate estimated power laws. *Inset:* distribution of E-I ratio. *Red:* low E-I ratios; *blue:* balanced E-I ratios; *gray*: high E-I ratios (highest 25% E-I ratios). **B)** Distribution of avalanche durations (*T*) for avalanches occurring at low (*red*) and balanced (*blue*) E-I ratios, respectively. Solid lines: estimated power laws. **C-D)** Same as (A) and (B) but comparing avalanches occurring at high E-I ratios (*gray*) and at balanced E-I ratios (*blue*). **E)** Power-law exponents describing the distributions of the sizes (*τ, top* panel) and durations (*α, bottom* panel) of neuronal avalanches occurring at low, balanced, and high E-I ratios. One-way ANOVA: for exponent *τ*: *F*_(2,27)_ = 7.35, *p* = 0.003; for exponent *α*: *F*_(2,27)_ = 7.94, *p* = 0.002. *: *p* < 0.05, post hoc t tests. Numbers in parentheses denote, for each E–I ratio category, the number of significant log-likelihood ratio tests supporting power-law behavior. **F-H)** Average power-law exponents *τ, α*, and *σνz* estimated for different bins of E-I ratios. Each E-I bin contained 1,000 neuronal avalanches, with the bin center corresponding to the mean E-I ratio of those avalanches. The relationship between exponents and E-I ratio was linearly interpolated and averaged across fish. Purple shaded areas indicate SEM, while gray lines and shaded areas show the expected critical exponents and their uncertainty in random field Ising models.

Finally, we also analyzed the proportion of excitatory and inhibitory activity within neuronal avalanches, i.e., the local E-I ratio, noted *k*. Unlike the global E-I ratio used for the previous analyzes, the local E-I ratio was calculated using only the activity of neurons participating in a given avalanche and therefore it was computed using nearby neurons. We found that neuronal avalanches involving only one cell type were small and short, whereas larger, longer avalanches exhibited a balanced proportion of E and I activity (**Figure 5A-C**). This suggests that neuronal avalanches propagated more when the recruited E and I activity of the participating neurons balanced each other. Moreover, we found that, on average, the size and duration of neuronal avalanches was maximal at a slightly excitatory-dominated local E-I ratio (*k*∼0.6) (**Figure 5D-E**). Then, we studied the average evolution of the size of neuronal avalanches of duration *T*, noted ⟨*S*(*t, T*)⟩ (**Figure 5F**), and the average temporal profiles of E and I activity during avalanches of duration *T*, denoted ⟨*S*_*E*_(*t, T*)⟩ and ⟨*S*_*I*_(*t, T*)⟩ (**Figure 5G**), respectively. As expected for systems operating near criticality, avalanche profiles collapse onto a single scaling function when time is normalized by the avalanche duration (*t*/*T*). This collapse is observed with comparable scaling exponents whether profiles are computed from the joint excitatory–inhibitory activity or separately for excitatory and inhibitory populations (see **Supplemental Note S2** and **Supplemental Figure S5**). Moreover, we found that during the avalanches, the local E-I ratio was slightly biased towards excitation. This slightly biased E-I ratio was maintained during the time-course of avalanches. Indeed, the ratio of E temporal profile to total activity during avalanches, defined as *ρ* = ⟨*S*_*E*_(*t, T*)⟩⁄(⟨*S*_*E*_(*t, T*)⟩ + ⟨*S*_*I*_(*t, T*)⟩), remained nearly constant, reflecting stable excitatory dominance during avalanches (**Figure 5H-I**). The latter indicates that excitatory-dominated E-I ratio was maintained during the time-course of avalanches, but with no evidence of a particular cell type participating more at their onset or termination.

**Figure 5.**
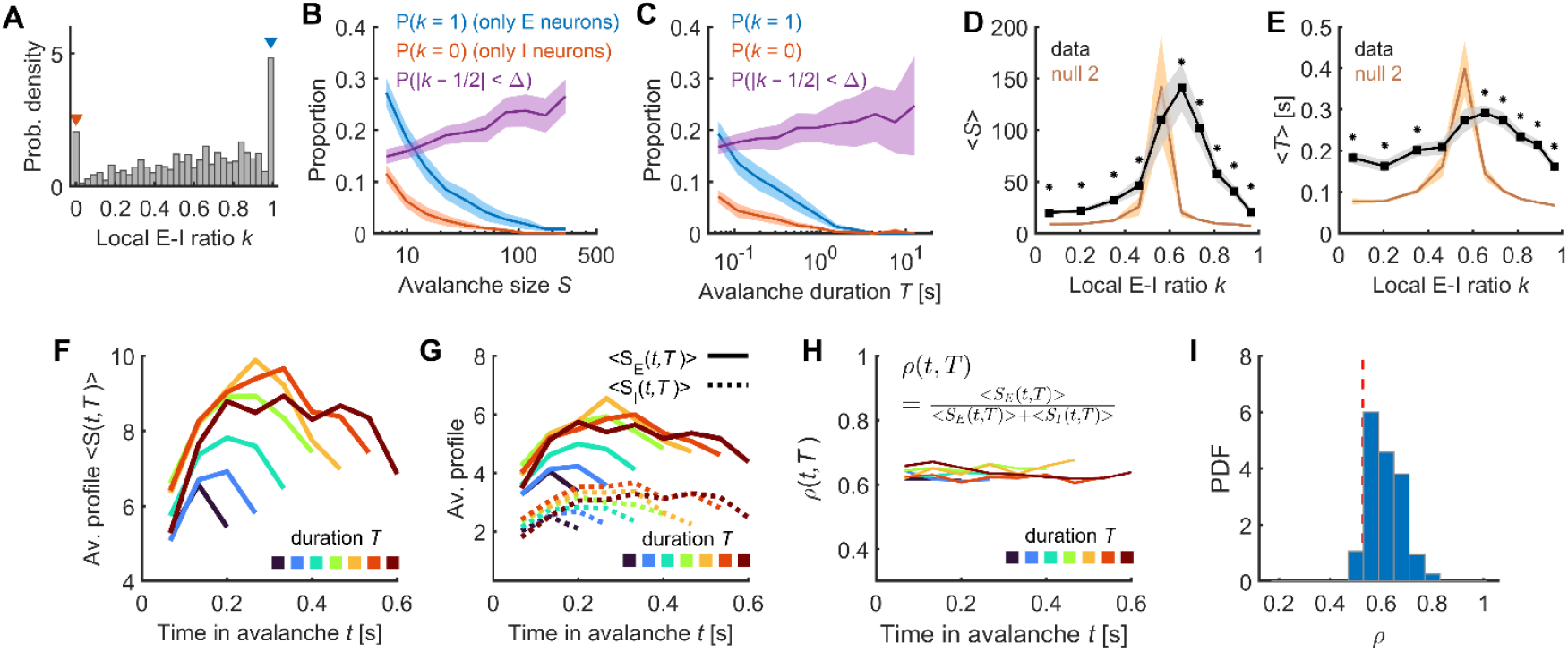
Local E-I ratio. **A)** Distribution of local E-I ratios, noted *k*, defined as the E-I ratio calculated using the activity of neurons participating in each neuronal avalanche. The *red* and *blue* arrowheads indicate avalanches composed entirely of inhibitory (*k* = 0) or excitatory (*k* = 1) neurons, respectively. **B-C)** Proportion of neuronal avalanches composed of only E (*blue*) or I neurons (*red*) as a function of avalanche size (B) and duration (C). The proportion of neuronal avalanches presenting a balanced local E-I ratio (|*k* − 0.5| < Δ, with Δ= 0.05) is shown in *purple*. Small and short avalanches were predominantly composed of a single cell type, whereas large and long avalanches exhibited a balanced E-I ratio. **D-E)** Average size (D) and duration (E) of neuronal avalanches across bins of local E-I ratios. The brown trace indicates the expected values of null models 2 (generated by permuting neuronal identities). Asterisks: significantly higher values with respect to null model 2 (*p* < 0.05). Significant differences from null model 2 indicate that the results depend on cell type rather than firing rates. **F)** Averaged temporal profile, ⟨*S*(*t, T*)⟩, of neuronal avalanches of durations *T* (data from fish #10). **G)** Averaged temporal profiles of E and I activity during avalanches of duration T, denoted ⟨*S*_*E*_(*t, T*)⟩ and ⟨*S*_*I*_(*t, T*)⟩, respectively (data from fish #10). **H)** Ratio of E temporal profile to total activity, *ρ* = ⟨*S*_*E*_(*t, T*)⟩⁄(⟨*S*_*E*_(*t, T*)⟩ + ⟨*S*_*I*_(*t, T*)⟩) (data shown for fish #10). **I)** Distribution of *ρ*, across all fish. The vertical red line indicates the average value of *ρ* expected given the firing rates of the E and I neurons.

### E-I network model suggests that neuronal avalanches emerge through balanced amplification

To study the neuronal mechanisms linking the E-I network ratio and critical dynamics of the optic tectum, we built a network model with E and I units capable of generating neuronal avalanches (see STAR **Methods**). To that end, we extended the stochastic Wilson-Cowan model^32,51^. In this model, E and I neurons interact through excitatory and inhibitory couplings, with strengths *w*_*E*_ and *w*_*I*_, respectively, and a nonlinear response function (**Figure 6A**). The neural network was built with spatial coordinates and cell-types given by the experiments. The neurons were connected with a probability that exponentially decreased with the distance between them (**Figure 6B-C**; see also STAR **Methods**). In order to compare between the model and the calcium imaging results, we used a convolution model^52^ to convert the model spiking activity into calcium-like transients like those observed in the experimental dataset (**Figure 6D**).

**Figure 6.**
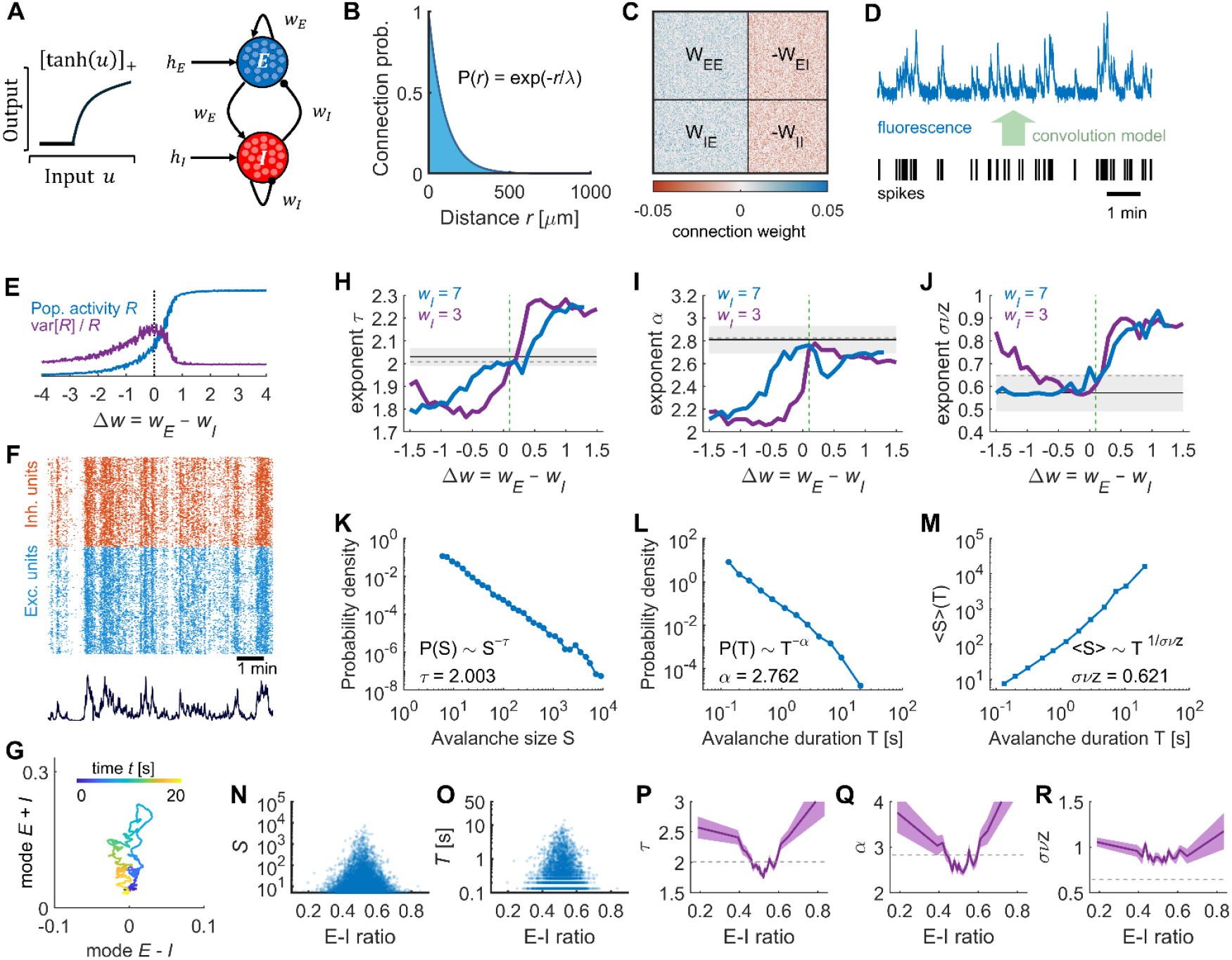
A stochastic Wilson-cowan model shows that neuronal avalanches emerge through balanced amplification. **A)** Model’s schematics (response function and network connectivity). **B)** Probability of connection as a function of distance. **C)** Connectivity matrix *W*, organized into submatrices *W*_*EI*_, *W*_*IE*_, *W*_*EE*_, and *W*_*II*_. **D)** A convolution model transformed the spiking activity into fluorescence signals, which were then thresholded to extract the calcium events. **E)** The population firing rate *R* increases a function of the difference between the excitatory and inhibitory coupling strengths, i.e., Δ*w* = *w*_*E*_ − *w*_*I*_, and its relative variance, *var* [*R*]⁄*R*, peaks when Δ*w* = 0.1. **F)** Example of model calcium events. *Top*: raster. *Bottom*: population calcium activity. **G)** Evolution of a neuronal avalanche in the activity space given by the difference (E−I) and sum modes (E+I). **H-J)** Power laws were fitted to the statistics of avalanches produced by the model’s calcium events, yielding estimated power-law exponents *τ* (G), *α* (H), and *σνz* (I) as a function of Δ*w* = *w*_*E*_ − *w*_*I*_, for varying *w*_*E*_ and *w*_*I*_ = 3 (purple trace) or for *w*_*I*_ = 7 (blue trace). The black horizontal line and the shaded area represent the expected critical exponents and their uncertainty, respectively. The dashed horizontal gray line indicates the average exponent value from the zebrafish data. The vertical dashed green line indicates Δ*w* = 0.1. **K-M)** Model’s probability distributions of avalanches sizes (K), durations (L), and their relation (M), for Δ*w* = 0.1 and *w*_*I*_ = 7 (see also **Supplemental Figure S7**). Estimated power-law exponents are indicated in the corresponding panels (see also **Supplemental Figure S8**). **N-R)** The sizes (N) and durations (O) avalanches of the model fluctuate spontaneously together with the network’s E-I ratio, leading to fluctuations in the avalanche exponents describing the distributions of sizes (P), durations (Q), and the relation between sizes and durations (R). Shaded areas indicate the estimation error of the exponents. Model parameters: Δ*w* = 0.1 and *w*_*I*_ = 7. Dashed horizontal line: average exponent value from the zebrafish data.

In the case of all-to-all connectivity, it has been shown that the stochastic Wilson-Cowan model has a critical point when the difference between the excitatory and inhibitory coupling strengths, i.e., Δ*w* = *w*_*E*_ − *w*_*I*_, reaches the critical value^51^ Δ*w*_*c*_ = 0.1. In this case, critical neuronal avalanches arise due to *balanced amplification*, a mechanism which amplifies the sum activity mode (E+I mode), while suppresses the difference activity mode (E−I mode)^32,53^. Balanced amplification is consistent with the experimentally observed strong positive correlation between the population activities of E and I neurons (**Figure 2E**). In the present sparse network, we also observed a phase transition around Δ*w*_*c*_ = 0.1, where the relative variance of the population spiking activity (before convolution) was maximized (**Figure 6E**)^51^. Around this point, the model produced calcium-like events resembling the empirical ones and exhibited evidence of neuronal avalanches (**Figure 6F**) evolving along the sum activity mode (**Figure 6G**; see also **Supplemental Figure S6**) as a result of strong amplification structure (BA index = 0.73; see STAR **Methods**). Specifically, the mean correlations between pairs of E-E, E-I, and I-I units were equal to 0.022, 0.020, and 0.018, respectively, closely matching the empirical values (see **Supplemental Figure S4B**). Notably, when analyzing the statistics of neuronal avalanches generated by the model’s calcium-like events for varying Δ*w*, we found that the power exponents {*τ, α, σνz*} matched the empirical ones in the region around Δ*w*_*c*_ = 0.1 (**Figure 6H-M**; see also **Supplemental Figures S7** and **S8**). This was also the case for the avalanche distributions calculated with the spatially unconstrained definition of neuronal avalanches (**Supplemental Figures S8B** and **S9A-F**). Moreover, the characteristics of neuronal avalanches produced by the model’s underlying spiking activity were consistent with previous reports using electrophysiological recordings^50^, including the associated power-law exponents and their behavior as a function of temporal coarse-graining (see **Supplemental Figure S9G-J**).

Finally, the neuronal avalanches produced by the model were smaller and shorter during fluctuating imbalances of the network’s E-I ratio (**Figure 6N-O**). Such fluctuations arise due to the stochastic nature of the model, but also due to the heterogeneous and sparse connectivity (**Supplemental Figure S10**). Consistent with our empirical observations, this led to deviations of the power-law exponents describing the neuronal avalanches (**Figure 6P-R**; see also **Supplemental Figure S11**), with exponents {*τ, α, σνz*} equal to {1.96, 2.65, 0.89} for E-I ratios between 0.4 and 0.6, {2.49, 3.39, 1.01} for E-I ratios lower than 0.4, and {2.63, 3.86, 1.02} for E-I ratios larger than 0.6. In conclusion, a stochastic Wilson-Cowan model at the critical balance between the excitatory and inhibitory coupling strengths was able to reproduce the statistics of neuronal avalanches observed experimentally and their dependence on the fluctuations of E-I ratios, suggesting that the observed critical activity in the optic tectum of the zebrafish could arise through balanced amplification.

## DISCUSSION

In this study, by monitoring the spontaneous dynamics of both excitatory and inhibitory neurons, we observed that the statistics of neuronal avalanches in the zebrafish tectum depended on the proportion of excitatory and inhibitory activity, as measured by the E-I ratio of the tectal network. We found that spontaneous neuronal avalanches exhibited critical-like dynamics at balanced E-I ratios but became smaller and shorter when the ratios were imbalanced, reflecting more ordered dynamics. Moreover, an extended stochastic network model of interacting E and I units, exhibiting a critical point for balanced excitatory and inhibitory couplings^51^, successfully reproduced the observed power-law statistics of zebrafish tectal neuronal avalanches. The model also predicted spiking activity avalanche exponents consistent with prior experimental findings^47–50^ and replicates the observed dependence of neuronal avalanches on E-I ratio fluctuations in the zebrafish tectum.

Extensive research further highlights the functional benefits of both E-I balance and critical dynamics. On one hand, balanced networks offer several advantages, including enhanced signal amplification, response selectivity, noise reduction, network stability, memory capacity, and facilitation of synaptic plasticity^53–59^. On the other hand, critical dynamics optimize neural computations by improving information transmission, storage, and processing^11,15–18,60^. Moreover, critical dynamics also emerge in complex environments^61,62^, in trained neural cultures^63^, and are restored following sleep^44^. Pioneering *in vitro* experiments have demonstrated that pharmacological manipulations of the E-I ratio shift neural circuits away from criticality, impairing their ability to process stimuli as quantified by dynamic range^36^, a measure that has been shown to peak at criticality in network models^64^. However, the link between criticality and E-I balance remains elusive. Here, by leveraging in vivo calcium imaging, immunostaining, and modelling, we demonstrated that the statistics of neuronal avalanches and their dependence on spontaneous E-I fluctuations in the zebrafish optic tectum align with a model that reaches criticality when excitation and inhibition synaptic couplings are strong and balance each other (**Figure 6F-R** and **Supplemental Figure S8**). In the present model, the neuronal avalanches arise from a mechanism known as *balanced amplification*. This mechanism promotes the joint activity of E and I units^32,53^ and provides a possible mechanism through which E-I balance leads to critical brain dynamics. Interestingly, previous theoretical work showed that balanced amplification can enhance signal propagation in neural networks^65^.

In this context, it is worth noting that previous studies have linked the E-I ratio to scale-invariant dynamics, with spiking network models suggesting that replicating experimental data requires 20–30% inhibitory synapses (as in the mammalian cortex) and stronger inhibitory synapses than excitatory ones^66,67^. In zebrafish larvae, however, the optic tectum contains similar numbers of excitatory and inhibitory cells with similar firing rates, indicating E-I balance through comparable synaptic strengths, which in the present model leads to a critical point. Consistent with this, the density of excitatory and inhibitory post-synaptic terminals has been shown to be similar in the optic tectum of zebrafish larvae^68^. Nevertheless, it is important to note that in our study the balance between excitation and inhibition activity is not a freely tunable control parameter (as the parameter *Δw* is in our model). Our experimental setup only allowed us to access the spontaneous activity of excitatory (E) and inhibitory (I) neurons, from which we tracked fluctuations of E activity relative to the total (E+I) activity. Excitation and inhibition were not experimentally manipulated, as in previous in vitro pharmacological studies; therefore, we cannot explore regimes in which either inhibition or excitation is selectively suppressed. Future studies could leverage optogenetic or chemogenetic approaches to directly control excitation and inhibition and systematically probe their role in shaping critical dynamics in vivo.

It is worth noting that our interpretation of the mechanisms underlying neuronal avalanches is inevitably influenced by the specific model adopted in this study. Alternative mechanisms can be envisioned. In particular, recent work has shown that slower synaptic dynamics of inhibitory neurons relative to excitatory neurons can drive networks of excitatory and inhibitory units to operate near a Hopf bifurcation—beyond which network oscillations emerge—and that, at this operating point, scale-invariant neuronal avalanches can arise together with several characteristic features of neuronal responses^21,22^. While such a mechanism may be relevant for mammalian cortex, it appears less likely in the case of zebrafish, as neuronal oscillations have not been reported in the zebrafish optic tectum. Nevertheless, future experimental and modeling work could explicitly test this alternative mechanism in the zebrafish brain.

Neuronal avalanches were identified using two previously established definitions: a spatially constrained method^8,11^, also applied in studies of magnetic domains^13^, and a spatially unconstrained approach^1–6,45^. We found that, while these definitions yielded different power-law exponents for the distributions of the zebrafish neuronal avalanche sizes, durations, and the size-duration relationship, both definitions produced exponents consistent with critical values predicted by the stochastic network model analyzed here and the spatially constrained definition yielded exponents consistent with those of the random-field Ising model^13,43,50^ (**Figure 1I, Figure 6K-M, Supplemental Figure S8**). In addition, with both definitions, the exponents also satisfy the crackling-noise relation. Taken together, these results reconcile the different approaches used to study neuronal avalanches and demonstrates their consistency with the criticality hypothesis. Indeed, the fact that the stochastic network model used here reproduces both sets of exponents at its critical point—depending solely on the avalanche definition employed—indicates that the two sets of exponents belong to the same universality class and differ only due to the method used to detect neuronal avalanches. A detailed comparison between the present stochastic Wilson–Cowan model and the random-field Ising model would require a fully dedicated study and is therefore beyond the scope of the current article.

Moreover, using the spatially constrained definition of avalanches, a local E-I ratio can be defined for each avalanche, and we observed that avalanches exhibited a slight excess of excitation (*k*∼0.6), while the global E-I ratio — which was calculated using all simultaneously recorded neurons— remained balanced (0.5). This suggests that inhibitory activity not participating in neuronal avalanches plays a compensatory role to counterbalance the excess of excitation within avalanches. However, this interpretation should be taken with caution, as inhibition alone may not fully explain why avalanches were smaller and shorter at high E–I ratios or even for avalanches composed exclusively of excitatory activity (Fig. 5B). This suggests that additional cellular and/or network mechanisms contribute to these effects and warrant further investigation. Finally, the spatially constrained definition of avalanches also enables the study of co-occurring avalanches^11^. Interestingly, recent theoretical work on directed networks has examined the co-occurrence of avalanches that do or do not share connections^69^. Future studies could adopt this approach by combining whole-brain data from zebrafish with zebrafish larva connectomes, which are currently under development.

Furthermore, the theoretical model allowed us to evaluate power-law exponents derived from different simulated signals, representing calcium events or spiking activity. For both signals, the model’s critical exponents align with experimental findings from calcium imaging and electrophysiology^11,47–50^. Also, at its critical point, the model replicates the effect of temporal coarse-graining on the power-law exponents of spiking neuronal avalanches, consistent with observations in electrophysiological recordings of the mouse cortex^50^ (**Supplemental Figure S9J**), and further supports the idea that these observations align with critical dynamics. Overall, our adapted theoretical model, at its critical point, reconciles key features of neuronal avalanches observed across different neural signals, temporal scales, species (mammals and vertebrates), and detection methodologies.

We note that our results are unlikely to be affected by spatial subsampling, which has been shown to introduce systematic biases when inferring collective neural properties^70^. In the present study, approximately half of the optic tectum was imaged. Importantly, we found very similar avalanche statistics to those reported in our previous work^11^, based on whole-brain recordings in which nearly all neurons of the larval brain were simultaneously monitored, effectively eliminating subsampling effects. Given the close agreement between these results and the present findings, we conclude that spatial subsampling is unlikely to play a significant role in our analysis.

Finally, an open question in neuroscience is how brain dynamics self-organize toward criticality. Criticality may emerge without fine-tuning through self-organizing mechanisms based on homeostatic plasticity rules^23–27^. Alternatively, in structurally heterogeneous networks, critical-like dynamics may extend over a range of parameters as Griffiths phases^71,72^, rather than being confined to a single critical point. Investigating the presence and properties of Griffiths phases in alternative models will require dedicated future work; a promising direction would be to incorporate heterogeneity in neuronal excitability into the present stochastic model.

In conclusion, our results show that E-I activity shapes the spontaneous neuronal avalanches in the zebrafish optic tectum in vivo. These results are consistent with a stochastic network model poised at a critical point with balanced excitation and inhibition couplings, suggesting that critical neuronal avalanches arise from balanced amplification.

## Acknowledgements

This study was supported by the Project PID2022-137708NB-I00 funded by MICIU/AEI /10.13039/501100011033 and FEDER, UE. A. Ponce-Alvarez was supported by a Ramón y Cajal fellowship (RYC2020-029117-I) funded by MICIU/AEI/10.13039/501100011033 and “ESF Investing in your future”, and by the Spanish State Research Agency, through the Severo Ochoa and María de Maeztu Program for Centers and Units of Excellence in R&D (CEX2020-001084-M). G. Sumbre and E.C.A. Hansen were supported by ERC CoG 726280. We thank the IBENS Imaging Facility for providing the platform for immunostaining imaging.

## Code availability

The codes to detect and analyze the neuronal avalanches and to model them are available at: https://github.com/adrianponce/ExcInh_Neuronal_Avalanches.

## Author contributions

Conceptualization, E.C.A.H., G.S., A.P.A.;

Formal analysis, M.J., M.A., E.C.A.H., S.N., V.C., A.P.A.;

Supervision, G.S., A.P.A.;

Funding acquisition, G.S., A.P.A.;

Methodology, E.C.A.H., G.S., A.P.A.;

Writing – original draft, M.J., M.A., G.S., A.P.A.;

Writing – review & editing, M.J., M.A., E.C.A.H., S.N., G.S., A.P.A.;

## Supplemental information

Document S1. Figures S1–S11, Tables S1-S2, Notes S1-2, and supplemental references.

## STAR METHODS

### KEY RESOURCES TABLE

| REAGENT or RESOURCE | SOURCE | IDENTIFIER |
| --- | --- | --- |
| **Experimental Models: Organisms/Strains** |  |  |
| Tg(HuC:H2B-GCaMP6f) | Dunn et al., 2016 | ZFIN: ZDB-ALT-150916-4 |
| Tg(Vglut2a-loxP-mcherry-loxP-gal4ff) | Satou et al. 2013 | ZFIN: ZDB-ALT-131127-2 |
| Tg(HuC:H2B-GCaMP6f) and Tg(Vglut2a-loxP-mcherry-loxP-gal4ff) | This paper | N/A |
| **Antibodies** |  |  |
| Rabbit Polyclonal Anti-Gad1b | GeneTex | GTX124460 |
| Goat Polyclonal Anti-Choline Acetyltransferase Antibody | Sigma-Aldrich | AB144P |
| Mouse Monoclonal Antibody to mCherry | EnCor Biotechnology Inc. | MCA-1C51 |
| Donkey anti-Rabbit IgG (H+L) Highly Cross-Adsorbed Secondary Antibody, Alexa Fluor™ Plus 647 | Invitrogen | A32795 |
| Donkey anti-Goat IgG (H+L) Cross-Adsorbed Secondary Antibody, Alexa Fluor™ 568 | Invitrogen | A11057 |
| Donkey anti-Mouse IgG (H+L) Highly Cross-Adsorbed Secondary Antibody, Alexa Fluor™ Plus 488 | Invitrogen | A32766 |
| DAPI | Sigma-Aldrich | D9542 |
| **Software and Algorithms** |  |  |
| µManager<br>(Image acquisition) | Edelstein et al. (2010) | https://micro-manager.org/ |
| MATLAB scripts (Data Analysis, stimulus control, scanner and piezo control) | This paper | https://www.mathworks.com/products/matlab.html |
| Ca2+ Imaging Toolbox | Romano et al. (2017) | doi.org/10.1371/journal.pcbi.1005526 |
| NCC MATLAB Toolbox | Marshall et al. (2016) | www.nicholastimme.com/software.html |
| Python package Powerlaw | Alstott et al. (2014) | https://github.com/jeffalstott/powerlaw |

#### Animal model

Zebrafish larvae were kept under a 14/10 hour on/off light cycle. They were raised in a 0.5x E3 embryo medium and, after 5 days post fertilisation (5dpf), were fed with paramecia. A double transgenic line was used, expressing Tg(HuC: H2B-GCaMP6f) in all neurons and Tg(Vglut2a-loxP-mcherry-loxP-gal4ff) in glutaminergic neurons. Calcium imaging experiments were conducted on 10 zebrafish larvae, 5-8 days post fertilisation. At this point no sex differentiation has yet occurred^73^, thus the sex of the animals is unknown. When under the microscope, larvae were embedded in 2% low-melting agarose and placed ventral side down on a coverslip platform (5mm high, 5 mm wide) within a chamber filled with embryo medium. As zebrafish larvae are translucent, this allowed for calcium imaging to be conducted in vivo, with no paralyzer agents or anaesthetics involved.

All experimental procedures were approved by the comiteé d’éthique en expérimentation animale n° 005. Reference number APAFIS#27495-2020100614519712 v14.2.2.

#### Selective-plane illumination microscopy

We recorded neural activity of the zebrafish optic tectum using selective-plane illumination microscopy (SPIM) at cellular resolution (see **Figure 1A** for area imaged). The calcium indicator (H2B-GCaMP6f) was excited by a micrometer-thick light sheet emitted from the left side of the larva which produced optical sectioning. A laser with wavelength of 488 nm (Omicron PHoXX® 480-200) was used as the single-photon light source of the SPIM. The laser beam was coupled to an optical single-mode fiber and the output was collimated using a fiber optic. The 1.6 mm beam from the coupled laser was expanded to a 4.5 mm beam through a telescope formed by two lenses (f1 AC254-050-A-ML and f2 AC254-150-A-ML, both from Thorlabs). The telescope created a collimated beam with uniform intensity. Next, the beam was projected onto a pair of galvanometric mirrors at an angular range of 26° (Model 6215H with a mirror size of 5mm). Then, a system composed of a scan lens (AC508 - 080 - AB -ML, Thorlabs) and a tube lens (Olympus U TLU IR) imaged the beam to a decentered spot at the back focal aperture of the imaging objective. The scan lens (fscan with a diameter of 50.8 mm) was placed in front of the scanning mirror and converted the angular deflection θ into a horizontal line displacement of the incident light. Next, the beam was collimated by an infinity corrected tube lens (ftube) and the beam was steered onto the back focal plane of the objective lens. The generated fluorescence of the sample was collected by a Hamamatsu ORCA-Flash4.0 V2® camera whose optical axis was orthogonal to the excitation plane. Overall, this created a stack of fluorescence images corresponding to three sections of the optic tecum for each zebrafish, recorded at 15Hz. The fluorescence images were recorded for a period of one hour, during which the fish was left in the dark, with as little auditory stimulation as possible.

#### Immunostaining

Immunostaining postmortem was used to distinguish between GABAergic, glutaminergic, and cholinergic neurons in the *in vivo* recording. The double transgenic (HuC:H2B-GCaMP6f and Vglut2a:loxP-mCherry-loxP-Gal4) zebrafish larvae were anesthetized and fixed in a solution of PBS+4% PFA at a pH of 7.4. Next, the fixed larvae underwent a series of washes in PBS+0.5% Triton X-100 to remove the fixative. A blocking solution containing PBS+0.5% Triton X-100, 1% DMSO, and 2% BSA was applied to saturate nonspecific binding sites. The samples were then incubated with primary antibodies diluted at 1/100 against specific markers of interest, including Rabbit Anti-Gad1b, Goat Anti-ChAT, and Mouse Anti-mCherry. After that, the samples were washed again and subjected to a secondary antibody staining step using Donkey Anti-rabbit 647, Donkey Anti-goat 568, and Donkey Anti-mouse 488 antibodies diluted at 1/500. After further washes, the samples were treated overnight with 500 µL of PBS, 1% Triton, 1% DMSO, 0.1% Tween, and DAPI (10 µg/µL) for nuclear staining. Finally, the samples were fixed once more with PBS+4% PFA, washed, and stored in PBS-0.5% solution at 4 ° C until confocal imaging. The samples were immobilized in 2% UltraPure™ Low Melting Point Agarose (16520100, Invitrogen™) on a glass-bottom µ-Dish 35 mm (high Glass Bottom) (81158, ibidi). Imaging was conducted using an inverted Confocal Laser Scanning Microscope TCS SP8 (Leica, Germany), equipped with a CS2 Plan Apochromat 40x/1.10NA water immersion objective lens and utilizing hybrid PMTs. To minimize any potential crosstalk between channels, images were acquired in frame sequential mode. All images were captured using a White Light Laser (WLL) with excitation wavelengths of 488 nm, 568 nm, and 647 nm to excite Alexa Fluor Plus 488, Alexa Fluor 568, Alexa Fluor Plus 647, respectively, as well as a 405 nm diode for DAPI excitation. The pinhole was set at 1.0 Airy unit. Imaging was performed with a 2 µ m Z-step interval at a resolution of 1024 x 1024 pixels. The zoom factor was 0.75. The immunostaining images were analyzed using the image processing package Fiji^74^. We aligned the images using the Fiji plugin BigWarp. Neurons that were not identified as ChAT or Vglut2a positive were considered as GABAergic.

#### Binary spontaneous neuronal activity data

For each fish, the spontaneous neuronal activity was recorded for a period of one hour, during which the fish was left in the dark, with as little auditory stimulation as possible. To detect calcium events, we binarized the activity of each of the *N* cells by thresholding the fluctuations of fluorescence intensity Δ*F*⁄*F* with a threshold equal to 3*σ*_*noise*_, where *σ*_*noise*_ is the standard deviation of the baseline fluctuations of the cell^75^. Above this threshold the activity was set to 1, otherwise it was set to 0. The temporal bin size was determined by the acquisition rate, equal to 15Hz, corresponding to a bin size of 66.67 ms.

#### Neuronal avalanches

Neuronal avalanches were defined following Ponce-Alvarez et al.^11^. At each time bin *t*, we detected clusters formed by nearby co-activated neurons. To each neuron, we associated a sphere of radius *R* = 10 μm, centered at 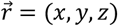, where 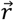 indicates the position of the neuron. Neurons were grouped to form clusters if their associated spheres overlapped and if they were active at the same time bin. A cluster is composed of at least 3 co-active, nearby neurons. At each time frame *t*, we obtained *m* clusters that we noted *C*_*i,t*_, where *i* ∈ {1, …, *m*}, with associated sizes (number of neurons) noted *C*_*s*_(*i*). Neuronal avalanches describe the spatiotemporal evolution of the activity clusters. A new avalanche was initiated at time *t*_*0*_ by the activation of a cluster 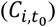 of neurons that were not active at the preceding time frame, i.e., 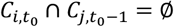. If at least one neuron of cluster 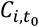 continued to be part of a cluster at time *t*_*0*_ + 1, i.e., 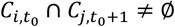, then the avalanche was continued, and so on, until this condition no longer held. An avalanche is described by its duration *T* (the time it lasts) and its size *S* (the number of neuronal activations during the avalanche). As shown previously^11^, with this definition, the statistics of neuronal avalanches were robust under coarse-graining in both time and space, in particular, for different choices of *R* (see also **Supplemental Figure S1C**). In addition, estimation of the influence radius *R*^∗^ from the decay of neuronal partial correlations as a function of inter-neuronal distance^76^ yielded an average radius ⟨*R*^∗^⟩ equal to 12.2 µm (see **Supplemental Figure S1D**).

The above definition of avalanches imposes a spatial constraint, allowing for the separation of two simultaneous avalanches occurring in distinct regions of the brain. This definition is different from that used in other studies on neuronal avalanches^1–6,45,46^. In those studies, neuronal avalanches were defined as consecutive time bins with at least one active site, regardless of whether activations formed spatially connected clusters. This non-spatial definition is applicable when studying activity within a specific brain region, as in our case (i.e., the optic tectum). According to this definition, the sum activity across all *N* recorded neurons is calculated at each time bin. An avalanche is then defined as a period during which this summed population activity exceeds a threshold *θ* (equal to 0.5% of the neurons). Neuronal avalanche statistics remained robust across a broad range of the threshold *θ* (**Supplemental Figure S3G**).

#### Power-law fitting

The probability distributions of durations and sizes of neuronal avalanches were approximated by truncated power-law distributions, *P*(*T*)∼*T*^−*α*^ and *P*(*S*)∼*S*^−*τ*^, using maximum likelihood estimation (MLE) as described in Marshall et al.^77^. The cutoffs used to truncate the data are indicated in **Tables S1 and S2**. The estimation errors of the power-law exponents were calculated using bootstrap re-sampling (1,000 re-samplings). The statistics of neuronal avalanches were compared to those obtained from surrogate datasets generated by randomizing the time indices of the *N*-dimensional activity vectors, a procedure that destroys temporal correlations while preserving correlations between neurons (**Supplemental Figures S1A-B** and **S3E-F**). The robustness of avalanche statistics with respect to the choice of truncation cutoffs in the power-law fits is shown in **Supplementary Figure S2**.

We tested the power-law hypothesis by determining whether it better describes the data compared to an alternative heavy-tailed distribution, specifically the lognormal distribution, by analyzing the log-likelihood ratio (LLR) between the two models. The lognormal distribution follows the density function: 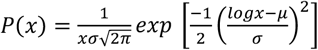, with dispersion parameter *σ* > 0 and location parameter *μ* > 0. For a given data x = (*x*_1_, …, *x*_*n*_), the LLR between the power-law and the lognormal was given by *LLR*(x) = *LL*_*PL*_ (x) − *LL*_*LN*_(x), where *LL*_*PL*_ and *LL*_*LN*_ are the log-likelihoods of the power law and the lognormal, respectively. If the likelihood of the power law model for a given empirical data set exceeds that of the lognormal model, the LLR is positive; otherwise, it is negative. To test whether the LLR is significantly different from zero, the p-value for the LLR test is given by: 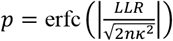, where erfc is the complementary error function, 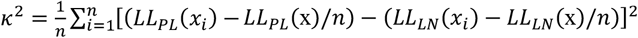, and 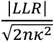 is the normalized log-likelihood ratio^78,79^. See **Tables S1 and S2**.

Since the above MLE approach applies only to probability distributions, it cannot be used to fit the relation between the average size ⟨*S*⟩(*T*) of avalanches of duration *T*. For this reason, to fit the power-law ⟨*S*⟩(*T*)∼*T*^−1⁄*σνz*^ we used least squares on log-log scattered data. We tested the power-law hypothesis by comparing the explained variance of the least-squares fit using the power law and the one obtained using an exponential function. Specifically, we calculated the ratio 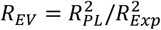, where 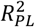 and 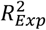 are the explained variances of the linear regression model log *Y* = *a* log *X* + *b* (power law) and log *Y* = *cX* + *d* (exponential), respectively. Ratios *R*_*EV*_ > 1 favor the power law hypothesis against the exponential alternative. The significance of *R*_*EV*_ > 1 was assessed using bootstrap re-sampling (5,000 re-samplings). The estimation error of the power-law exponent was given by the error of the slope of the linear regression model. See **Tables S1 and S2**.

#### Network model

We modelled the collective activity of E and I neurons using a model of critical dynamics that combines stochastic Wilson-Cowan equations^32,51^ and spatial embedded neuronal connectivity. Briefly, this model describes the dynamics of *N*_*E*_ excitatory neurons and *N*_*I*_ inhibitory neurons. A neuron *i* can be in an active (*s*_*i*_ = 1) or inactive state (*s*_*i*_ = 0) and evolves according to a master equation governing the two-state Markov process. The transition rate from an active state to a quiescent state is *ρ* for all neurons, i.e., *P*(1 → 0,in time *dt*) = *ρdt* (here *ρ* = 0.01 s^-1^). The transition rate from quiescence to activation depends on neuron’s inputs through the activation function: *P*(0 → 1,in time *dt*) = *f*(*u*_*i*_)*dt*, for neuron *i*, where *f* is the input-output (or transfer) function of the neurons, defined as *f*(*u*) = [*tanh*(*u*)]_+_. Here, 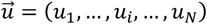 represents the synaptic input defined as: 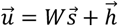, where 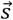 and 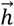 are *N*-dimensional vectors (*N* = *N*_*E*_ + *N*_*I*_) representing the state of the network and the external input to it, respectively. *W* represents the connectivity matrix coupling the *N* neurons. Here, we set *N*_*E*_ and *N*_*I*_ equal to the number of recorded excitatory and inhibitory neurons in the different SPIM experiments.

We also assigned spatial coordinates and cell types to the model neurons based on the positions and cell types of the recorded excitatory and inhibitory neurons. Additionally, we assumed that the connection probability, *p*_*ij*_, between two neurons decreases exponentially with the distance between them, i.e., 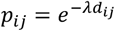, where *d*_*ij*_ is the Euclidean distance between neurons *i* and *j* given by the SPIM data. This resulted in an adjacency matrix *A* for which *A*_*ij*_ = 1 with probability *p*_*ij*_. We imposed *A*_*ii*_ = 0 (i.e., no self-coupling). The parameter *λ* determines the connectivity decay as a function of distance, controlling the spatial extent of the connectivity. thereby controlling the spatial extent of the connections. This exponential distance rule (EDR) has been reported in prior studies using retrograde fiber tracing methods in mammalian cortices^80^ and has been proposed as a simple, geometry-constrained wiring principle^81,82^. The connections that a given excitatory neuron *i* makes with the rest of the neurons were given as *W*_*ij*_ = *A*_*ij*_ *w*_*E*_⁄*K*_*i*_, where *K*_*i*_ is a normalization factor such that all excitatory neurons have the same connectivity strength *w*_*E*_, i.e., *K*_*i*_ = ∑_*j*_ *A*_*ij*_. Analogously, the connections that a given inhibitory neuron *i* makes with the rest of the neurons were given as *W*_*ij*_ = −*A*_*ij*_ *w*_*I*_⁄*K*_*i*_, where −*w*_*I*_ is the connectivity strength of inhibitory neurons.

In the case of all-to-all connectivity and vanishing external input *h*, it has been shown^51^ that the model presents a critical point when the difference between the excitatory and inhibitory coupling strengths, i.e., Δ*w* = *w*_*E*_ − *w*_*I*_, reaches a critical value Δ*w*_*c*_ equal to Δ*w*_*c*_ = *ρ* = 0.1. At this point, critical avalanches are observed due to strong balanced amplification^32,51,53^. Indeed, in the all-to-all connectivity case, where a mean-field approximation is applicable, it can be shown that the difference mode (*r*_*E*_ − *r*_*I*_, where *r*_*E*_ and *r*_*I*_ are the proportions of active E and I units) always has a fixed point at zero (i.e., *r*_*E*_ = *r*_*I*_)^32^. Fluctuations around this fixed point can then drive the sum mode (*r*_*E*_ + *r*_*I*_) due to the feedforward structure that appears when the dynamics are expressed in the sum–difference basis, resulting in neuronal avalanches^32^. For any connectivity *W*, a balanced amplification (BA) index can be defined to quantify the strength of hidden feedforward structure in the network^53^. This index measures the degree to which the dynamics of different eigenmodes are driven by feedforward interactions. The BA index is defined as the sum of the squared magnitudes of the feedforward (off-diagonal) components in a Schur decomposition of **W**, divided by the sum of the absolute squares of all of the elements: 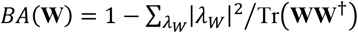, where *λ*_*W*_ are the eigenvalues of **W**, and Tr is the trace, and **W**^†^ is the conjugate transpose of **W**. A BA index equal to 0.70 indicates that 70% of the total power in the matrix driving the dynamics is in the feedforward links.

The network’s dynamics can be simulated as a continuous-time Markov process, using Gillespie’s algorithm^83^. Specifically, first, for each neuron *i* the transition rate *r*_*i*_ is computed, with *r*_*i*_ = *ρ* or *r*_*i*_ = *f*(*u*_*i*_) = *f*(∑_*j*_ *W*_*ij*_*s*_*j*_ + *h*_*i*_) if neuron *i* is active or quiescent, respectively; second, the network’s transition rate is calculated as *r* = ∑_*i*_ *r*_*i*_; third, the time interval *dt* is drawn from an exponential distribution with rate parameter *r*; finally, the state of neuron *i* is updated with probability *r*_*i*_⁄*r* and time is updated to *t* + *dt*. The network activity was simulated for a time corresponding to 30 min. Using numerical simulations, we found that the relative variance of population spiking activity peaks around Δw ≈ 0.1 (Fig. 6E; peak at Δw = 0.08 ± 0.01), consistent with analytical results for all-to-all connectivity, where activity variance is maximal at the critical point^51^. Note that the model’s avalanche behavior is similar under both all-to-all and sparse connectivity^32^.

An analytical determination of the critical point is only feasible for all-to-all connectivity (de Candia et al.). However, since avalanche behavior is similar under all-to-all and sparse connectivity (Benayoun et al.), we estimate the critical point numerically by locating the peak of the relative variance of population spiking activity, yielding

The model’s spiking activity was transformed into fluorescence signals through a convolution model^52^. This phenomenological model converts spike times to synthetic fluorescence time series, using a double-exponential kernel and a nonlinear transformation. Briefly, let {*t*_*k*_} be the set of spikes generated by a given model neuron, a latent variable *c*(*t*) results from the deterministic convolution of the spikes, *c*_*D*_(*t*), plus a noise term, as: 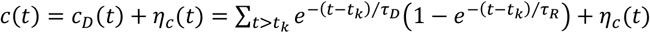, where *τ*_*R*_ and *τ*_*D*_ represent the rise and decay times of the calcium indicator, and *η*_*c*_ is Gaussian “internal” noise (equal to 0.1× ⟨*c*_*D*_⟩, i.e., 10% of the average deterministic term). We chose *τ*_*R*_ = 0.5 s and *τ*_*D*_ = 3 s, corresponding to the rise and decay times of the calcium indicator used in the experiments^84,85^. The latent variable *c*(*t*) was then converted to a synthetic fluorescence signal through a sigmoidal function^52^: *Y*_fluo_(*t*) = *σ*[*c*(*t*)] + *η*_*e*_(*t*), where *η*_*e*_(*t*) represents external noise (equal to 0.1× ⟨*σ*[*c*(*t*)]⟩) and *σ*[*c*(*t*)] = *F*_*m*_[1 + *e*^−*K*(*c*(*t*)−*κ*)^]^−1^, with *F*_*m*_ = 10, *K* = 0.6, and *κ* = 5. The fluorescence signals were downsampled at the resolution of the present experiments, i.e., 15 Hz. Finally, we binarized the synthetic fluorescence signal of each of the *N* cells by thresholding *Y*_fluo_(*t*) with a threshold equal to three standard deviations.

The model results are shown in **Figure 6**, where the model was informed by the spatial coordinates and cell types recorded in zebrafish larva #3. Results are equivalent using data from all zebrafish larvae (see **Supplemental Figure S8**).

## SUPPLEMENTAL INFORMATION

### Supplemental figures

**Supplemental figure S1.**
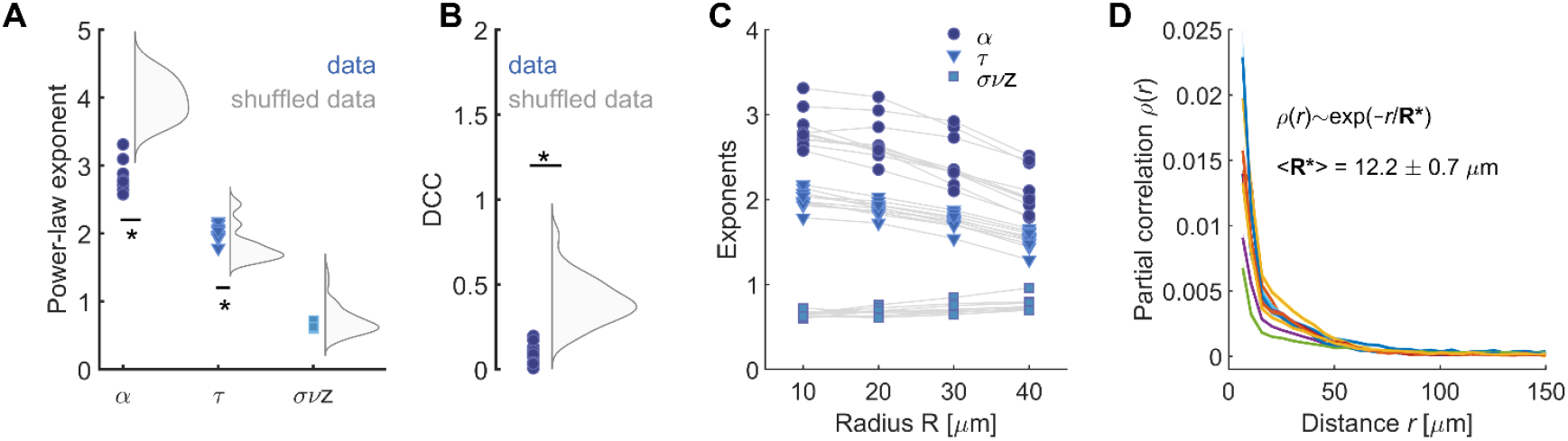
Controls on neuronal avalanche statistics. **A)** Comparison of power-law exponents obtained from the data with those obtained from surrogate datasets generated by randomizing the time indices of the *N*-dimensional activity vectors. This procedure destroys temporal correlations while preserving spatial correlations (10 surrogates per fish). *: p<0.001 (*t*-test comparing data exponents with the surrogate distributions). **B)** DCC values computed from the data and compared with the distribution of DCC values obtained from surrogate datasets. *: p<0.001 (*t*-test). **C)** Dependence of the estimated power-law exponents on the radius *R* used to detect neuronal avalanches. Consistent with Ponce-Alvarez et al.^11^, the exponents remained stable for radii up to ∼30 µm. **D)** Partial correlation functions for the different fish. Partial correlation function *ρ*(*r*) calculated as a function of the distance *r* between neurons (color traces correspond to *ρ*(*r*) for each fish). The characteristic radius *R*^*^ was then estimated by fitting *ρ*(*r*) with an exponential decay function, i.e., *f*(*r*) = *K* exp(−*r*/*R*^∗^). The resulting average radius ⟨*R*^∗^⟩ was equal to 12.2 µm, in good agreement with the value used to detect the neuronal avalanches (*R* = 10 µm).

**Supplemental figure S2.**
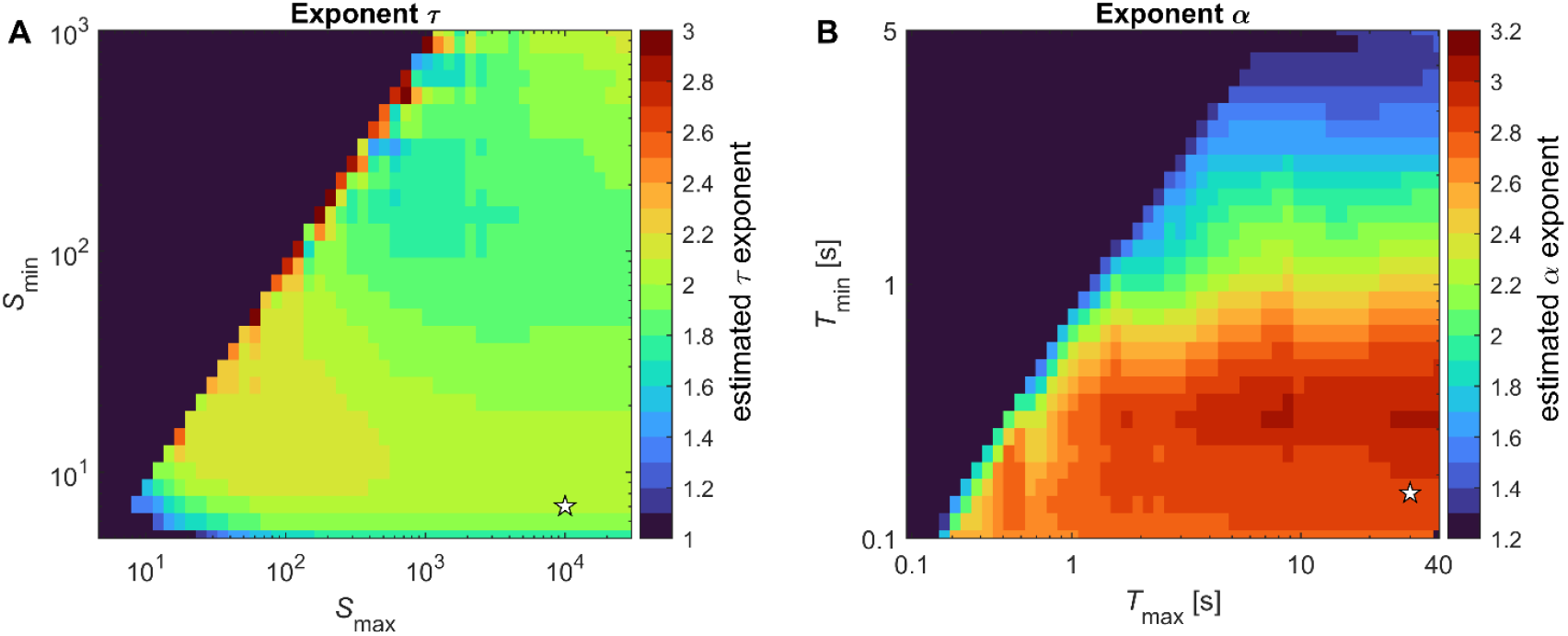
Dependence of power-law exponents on truncation cutoffs. **A-B)** Average exponents τ (A) and α (B) as a function of truncation cutoffs [*S*_min_; *S*_max_] and [*T*_min_; *T*_max_], respectively. Exponents were estimated using MLE. The stars indicate the cutoffs used in Table S1. Note that the estimated exponents remain stable over a wide range of cutoff values (provided that the upper cutoff sufficiently exceeds the lower cutoff).

**Supplemental figure S3.**
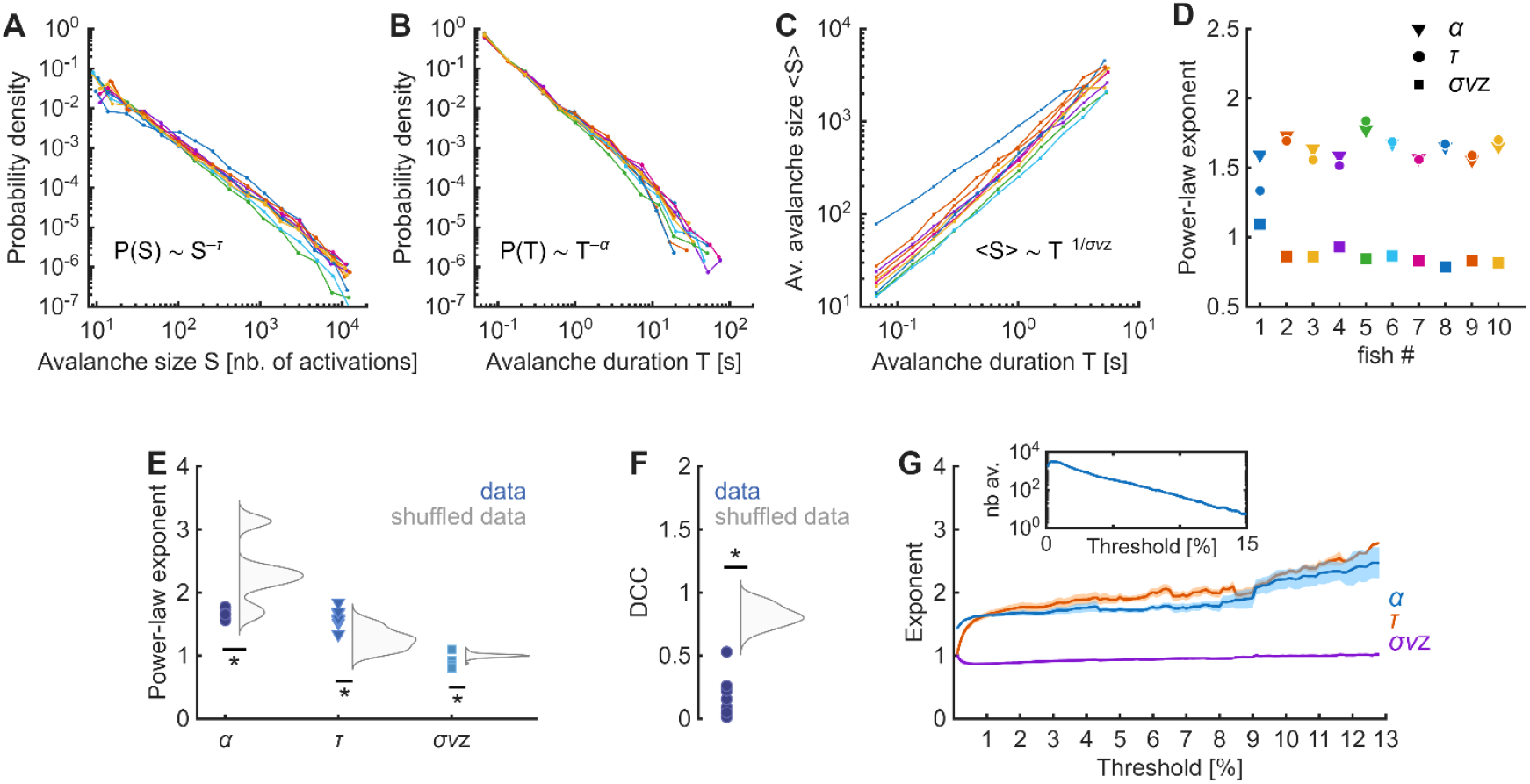
Neuronal avalanches using the spatially unconstrained avalanche definition. **A)** Distribution of avalanche sizes *S*. **B)** Distribution of avalanche durations *T* (in seconds). **C:** Relation between ⟨*S*⟩ and *T*, for each fish. In (A), (B), and (C) each color corresponds to a fish. The validity of the power-law fitting was evaluated using log-likelihood ratio tests. **D)** Measured exponents for each fish. Triangles: *α* exponent; circles: *τ* exponent; squares: *σνz* exponent. Error bars (estimation errors) are smaller than the size of the symbols. Individual power-law exponents, estimation errors, and power-law tests are detailed in **Table S2. E)** Comparison of power-law exponents obtained from the data with those obtained from surrogate datasets generated by randomizing the time indices of the N-dimensional activity vectors. This procedure destroys temporal correlations while preserving spatial correlations (10 surrogates per fish). *: p<0.001 (t-test comparing data exponents with the surrogate distributions). **F)** DCC values computed from the data and compared with the distribution of DCC values obtained from surrogate datasets. *: p<0.001 (t-test). **G)** Dependence of the estimated power-law exponents on the activity threshold used to detect neuronal avalanches. Exponents are stable for thresholds between 0.5 and 9%. Inset: number of detected avalanches as a function of the threshold. Note that for thresholds above 15%, the number of avalanches drops below 100.

**Supplemental figure S4.**
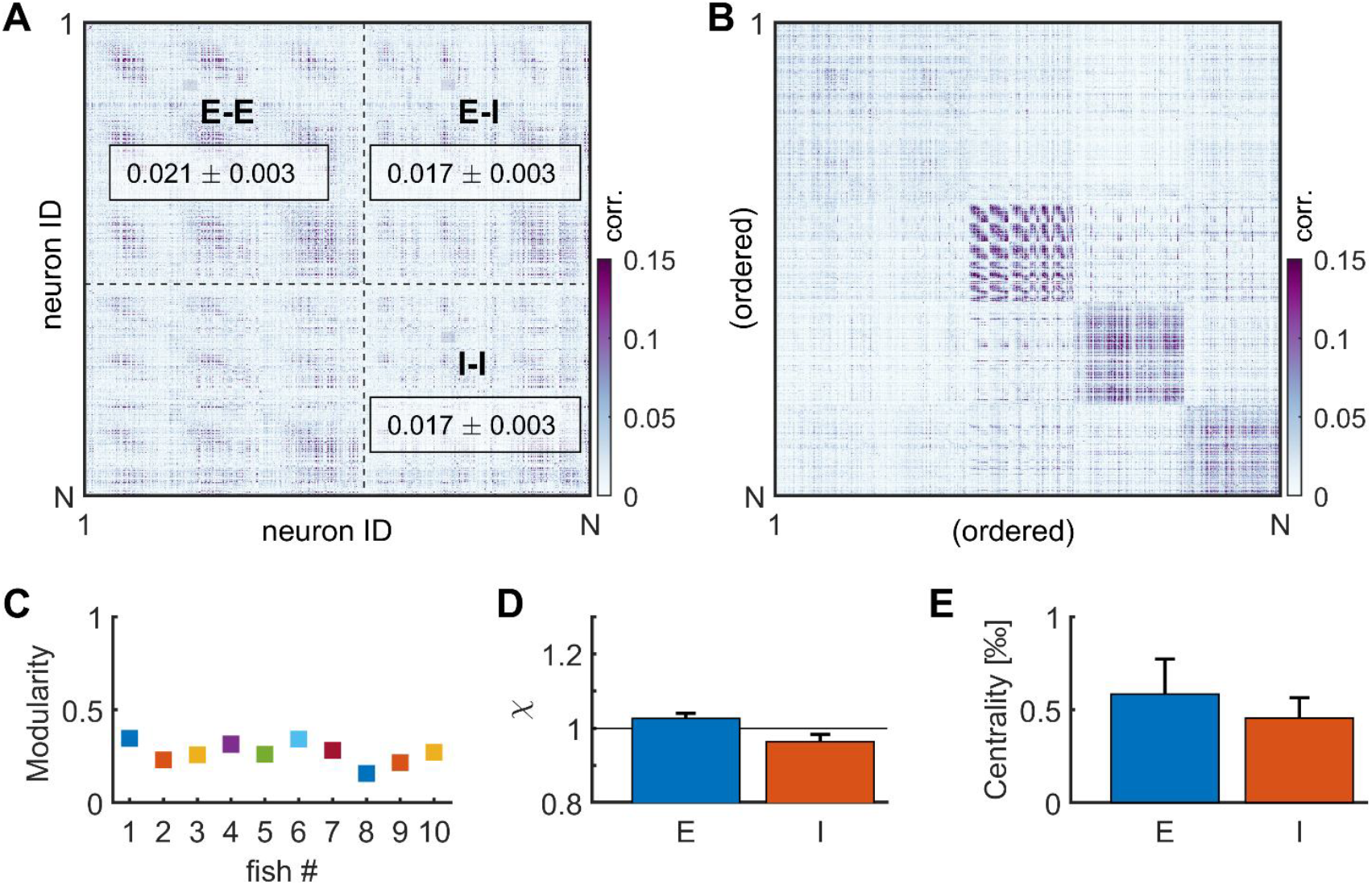
Stationary activity correlations. **A)** Correlation matrix among recorded neurons of an example fish. The matrix was ordered according to cell type. The boxes indicate the correlations among E neurons, among I neurons, and between E and I neurons, averaged across all corresponding pairs of neurons and all zebrafish larvae. **B)** Here the correlation matrix was ordered according to module participation (data from the same example fish as in panel A). **C)** Modularity index of the correlation matrix, for each fish. **D)** For each module, we calculated the ratio *χ* as the proportion of type *j* neurons in the module relative to the proportion expected in randomly selected groups of cells of the same size as the module. A value of *χ* near 1 indicates that the module was not significantly enriched for a specific cell type. Mean *χ* values are indicated for each cell type. Error bars indicate SEM. **E)** Average betweenness centrality of neurons of different type. Error bars indicate SEM.

**Supplemental figure S5.**
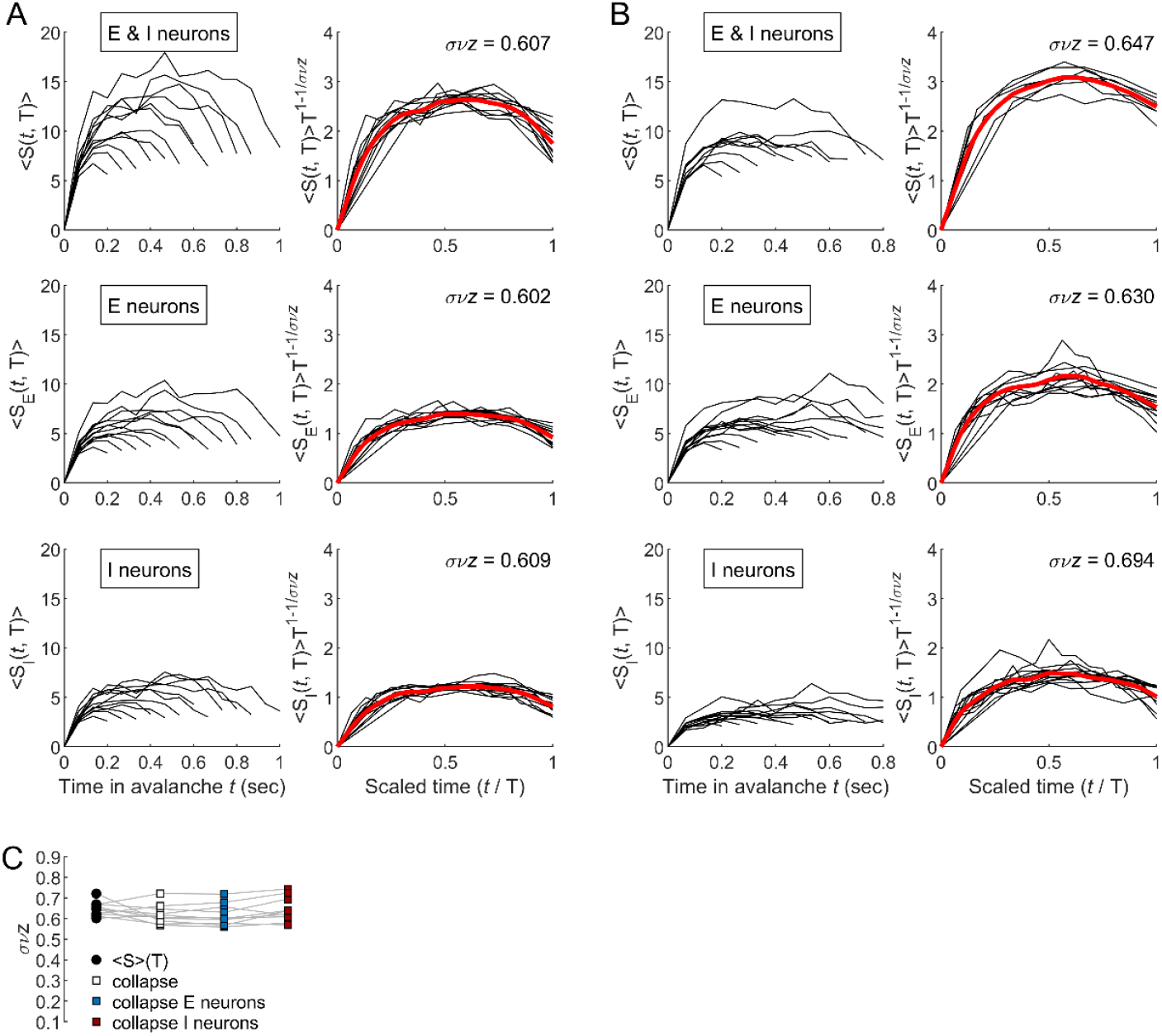
Shape collapse of neuronal avalanches. Close to criticality, the average avalanche profile, is expected to be similar across temporal scales (see Supplemental Note S2). **A) *Top*:** *Left:* Averaged temporal profile, ⟨*S*(*t, T*)⟩, of avalanches of durations *T* (data from fish #4). *Right:* Scaled avalanche profiles as a function of the scaled time *t/T*. Red line: averaged scaled avalanche profile; *σνz*: best scaling parameter. **Middle:** *Left:* Temporal profiles of E activity during avalanches of duration *T*, denoted ⟨*S*_*E*_(*t, T*)⟩. *Right:* Scaled avalanche profiles of E activity. **Bottom:** *Left:* Temporal profiles of I activity during avalanches of duration *T*, denoted ⟨*S*_*I*_(*t, T*)⟩. *Right:* Scaled avalanche profiles of I activity. **B)** Same as (A) but for fish #10. **C)** Estimated *σνz* exponents using the relation ⟨*S*⟩(*T*) (circles) and scaling collapse (squares). Each marker represents a different fish. Note the similarity between the exponents calculated with the two different methods and using the different types of avalanche profiles.

**Supplemental figure S6.**
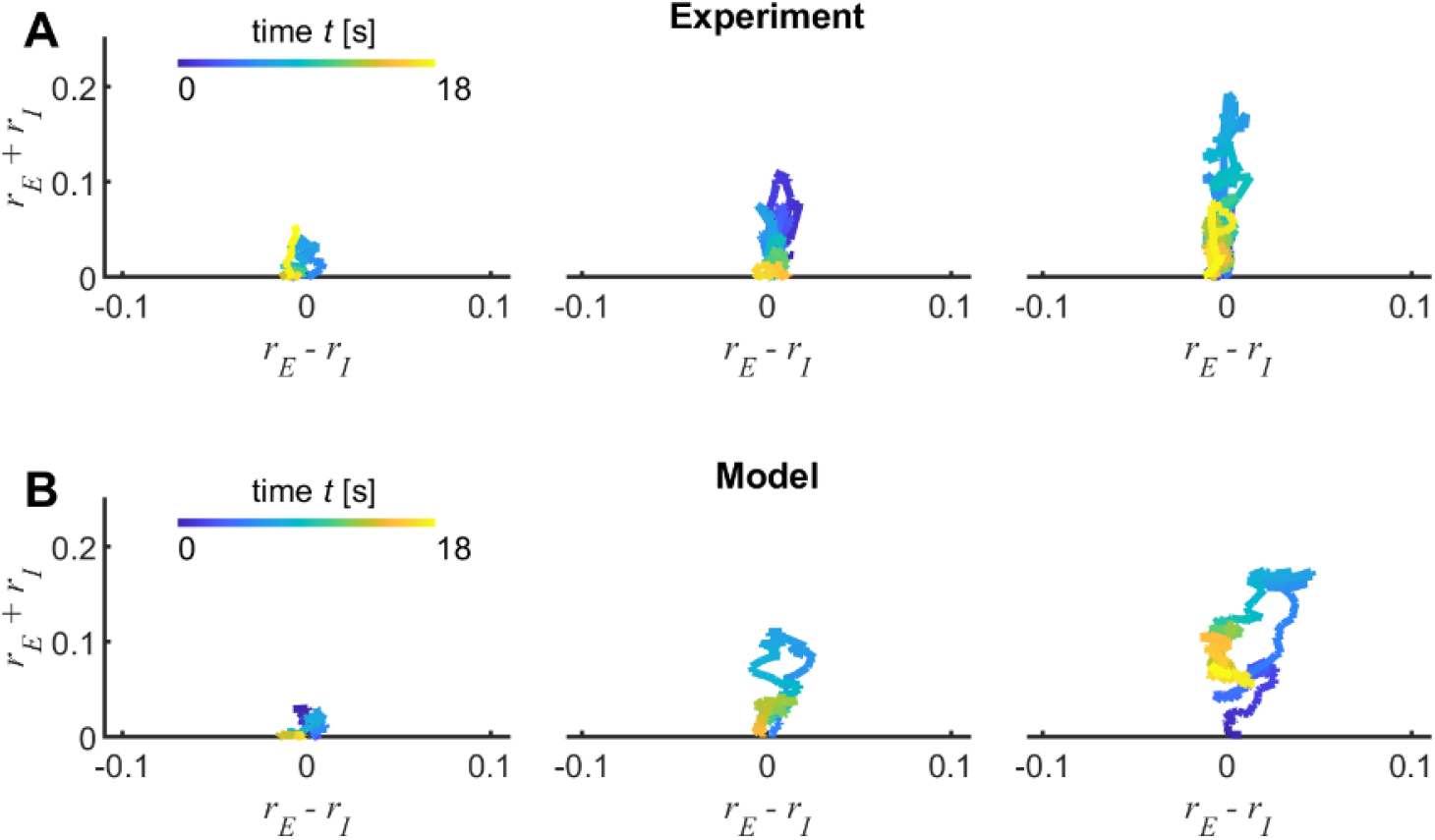
Difference and sum modes during neuronal avalanche activity. Evolution of neuronal avalanche in the activity space given by the difference (E−I) and sum modes (E+I), shown for one example fish (A) and for the model at its critical point (B). Three avalanches of different sizes are presented for both the experimental data and the model.

**Supplemental figure S7.**
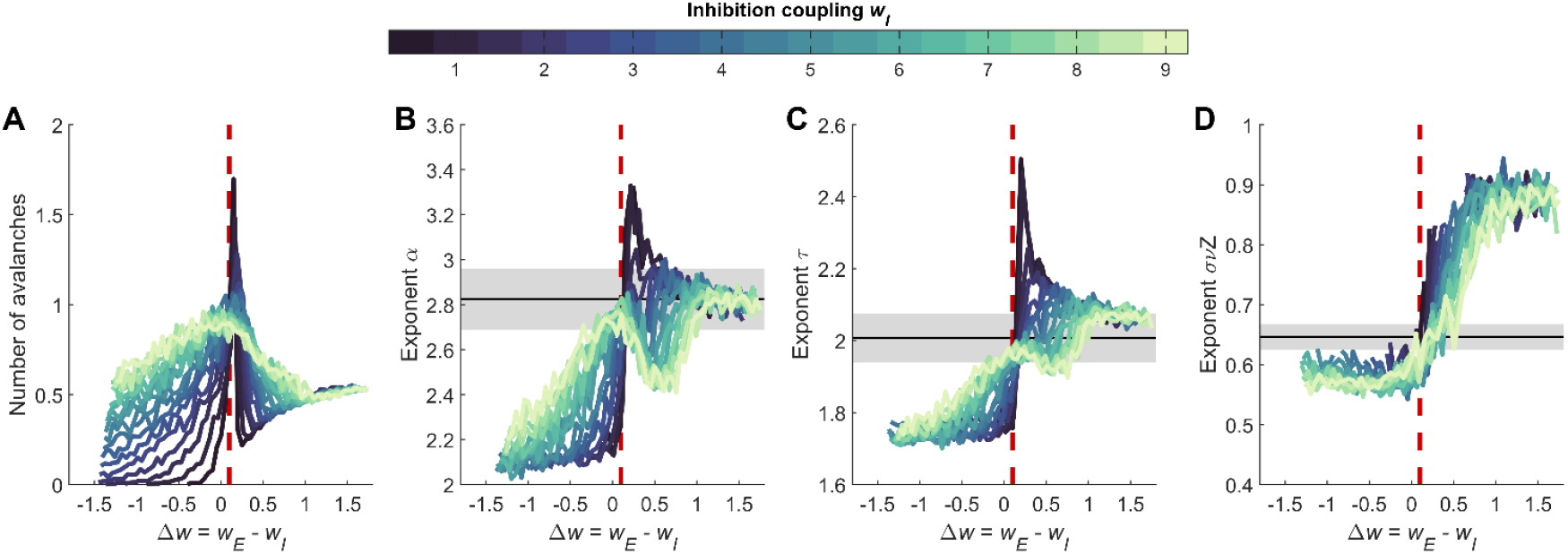
Model avalanche statistics as a function of *Δw* and *w*_*I*_. Neuronal avalanche statistics generated by the model were analyzed as a function of the difference between excitatory and inhibitory couplings, *Δw* = *w*_*E*_ − *w*_*I*_, while keeping *w*_*I*_ fixed. For each pair of values (*w*_*E*_, *w*_*I*_), network activity was simulated for a duration equivalent to 30 minutes. **A)** Number of neuronal avalanches. **B)** Estimated exponent α of the avalanche duration distribution. **C)** Estimated exponent τ of the avalanche size distribution. **D)** Estimated exponent σνz characterizing the scaling relationship between average avalanche size and duration. In panels (B)–(D), vertical red lines indicate *Δw* = 0.1, while the horizontal black line and gray shaded area denote the mean and SEM of the corresponding exponents measured in the zebrafish data. Note that for low *w*_*I*_, the parameter region in which the model fits the data is narrower than for larger *w*_*I*_.

**Supplemental figure S8.**
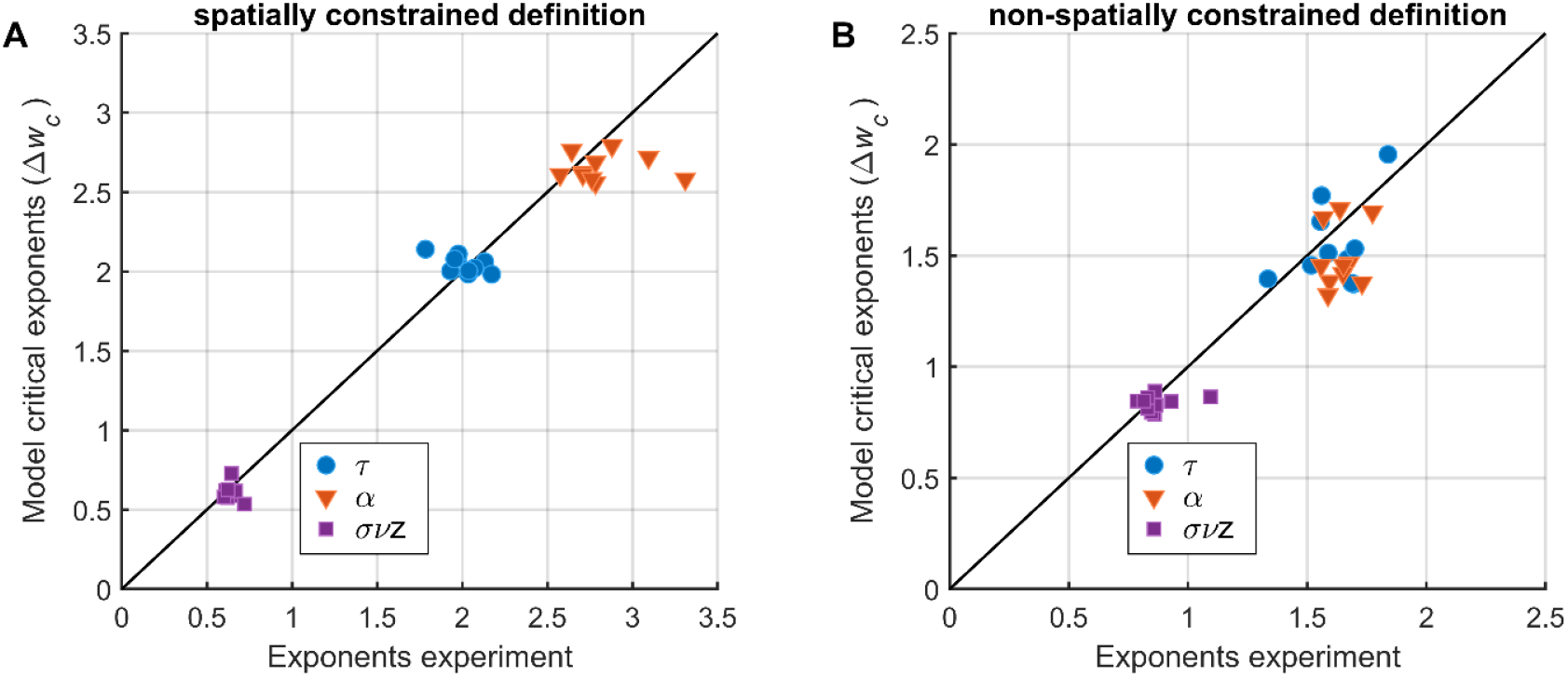
Neuronal avalanches of the model at its critical point. The model’s connectivity was constructed based on experimental data for each zebrafish. Specifically, the numbers of excitatory and inhibitory units were matched to the corresponding numbers of recorded neurons in the SPIM experiments. Model neurons were assigned spatial coordinates and cell types as determined by the experiments, and their connections were established with probabilities that decreased exponentially with distance. Avalanche analysis was performed on the simulated calcium activity using the spatially (A) and non-spatially (B) constrained avalanche definitions, respectively. The power exponents {*τ, α, σνz*} of the neuronal avalanches were calculated at the model’s critical point, i.e., Δ*w*_*c*_ = 0.1. These exponents were compared to those measure in the data for each fish.

**Supplemental figure S9.**
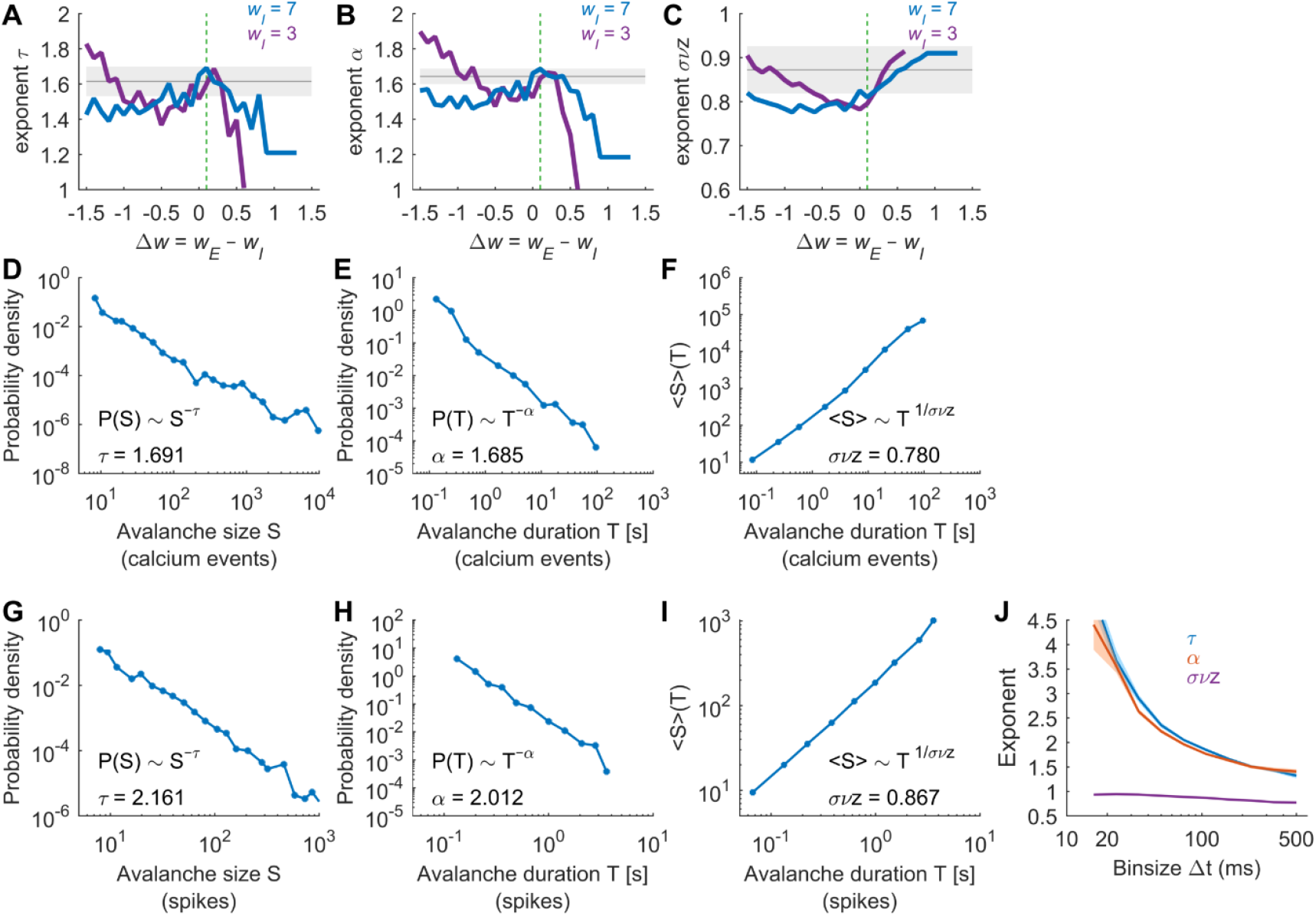
Neuronal avalanches of the model’s calcium transients and spiking activity, calculated using the spatially unconstrained avalanche definition. **A-C)** Power-law exponents *τ* (A), *α* (B), and *σνz* (C) of the model as a function of Δ*w* = *w*_*E*_ − *w*_*I*_, for *w*_*I*_ = 3 and varying *w*_*E*_ (purple trace) and for *w*_*I*_ = 7 and varying *w*_*E*_ (blue trace). Neuronal avalanches were calculated using the model’s calcium transients and the spatially unconstrained avalanche definition. The gray horizontal line and shaded area represent the average empirical exponents and their uncertainty, respectively. The vertical dashed green line indicates Δ*w* = 0.1. **D-F)** Probability distribution of the model’s avalanche sizes (D), avalanche durations (E), and the relationship between average size and duration (F) for model calcium events with Δ*w* = 0.1 and *w*_*I*_ = 7. **G-I)** Same as (D-F), but for the model’s spiking activity. As with calcium events, neuronal avalanches were calculated using time bins of Δ*t* = 66.67 ms (matching the resolution of the SPIM experiments, i.e., 15 Hz). **J)** The behavior of the avalanche exponents for spiking activity as a function of Δ*t* aligns with previous reports using *in vivo* spiking activity^50^.

**Supplemental figure S10.**
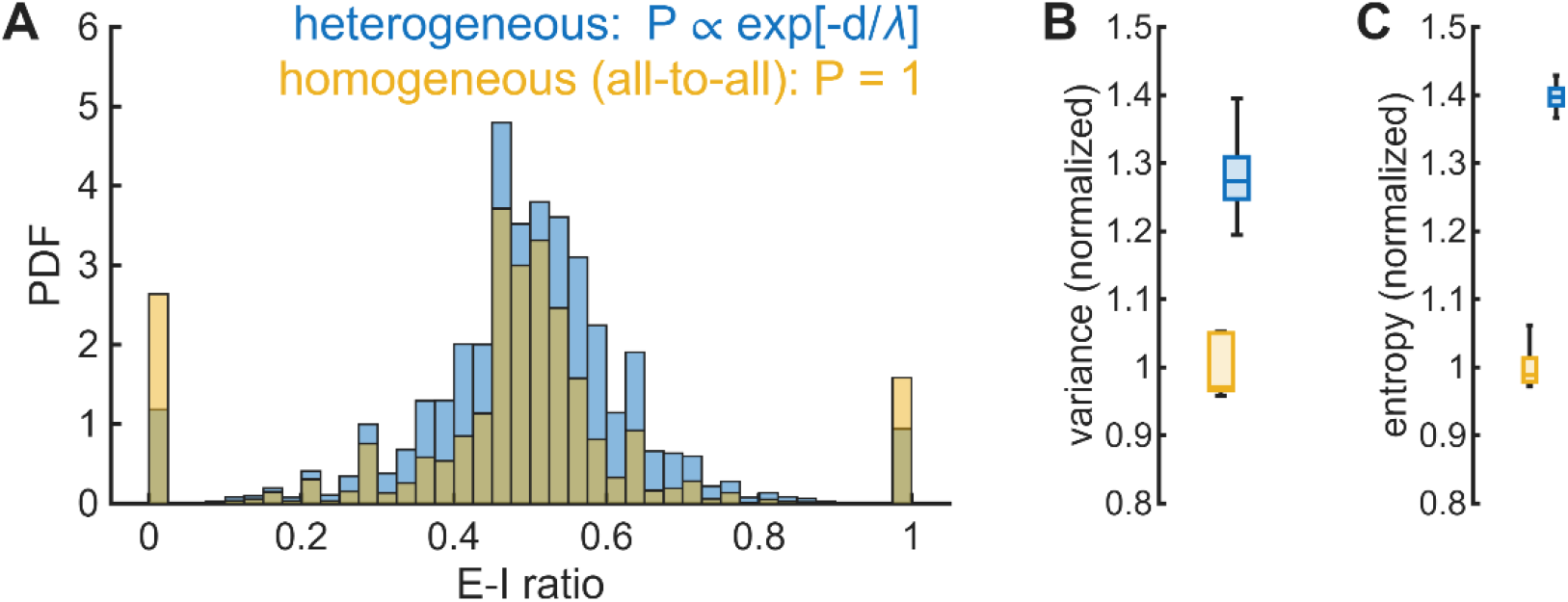
Fluctuations of E-I ratio for homogeneous and heterogeneous models. **A)** We compared the distribution of E-I ratios for the original model, with heterogeneous connections resulting from the exponentially-decaying connection probability, and for the homogeneous model for which connectivity is all-to-all (i.e., connection probability equal to 1). For both homogeneous and heterogeneous models we set Δ*w* = 0.1. Ten stochastic realizations were used for homogeneous and heterogeneous models, respectively. In the homogeneous model, fluctuations of the E–I ratio arise solely from the stochastic activation of the units, whereas in the heterogeneous model, additional fluctuations come from the variability in connectivity across units. **B)** Variance of E-I ratios were calculated for non-extreme values (i.e., E-I ratios different than 0 or 1). The variance was normalized to the mean variance of homogeneous model. **C)** The entropy of E-I ratios was calculated for the entire distribution of E-I ratios. The entropy was normalized to the mean entropy of homogeneous model. Panels (B) and (C) indicate that heterogeneous connectivity leads to a 30–40% increase in the variability of the E–I ratio relative to the homogeneous model.

**Supplemental figure S11.**
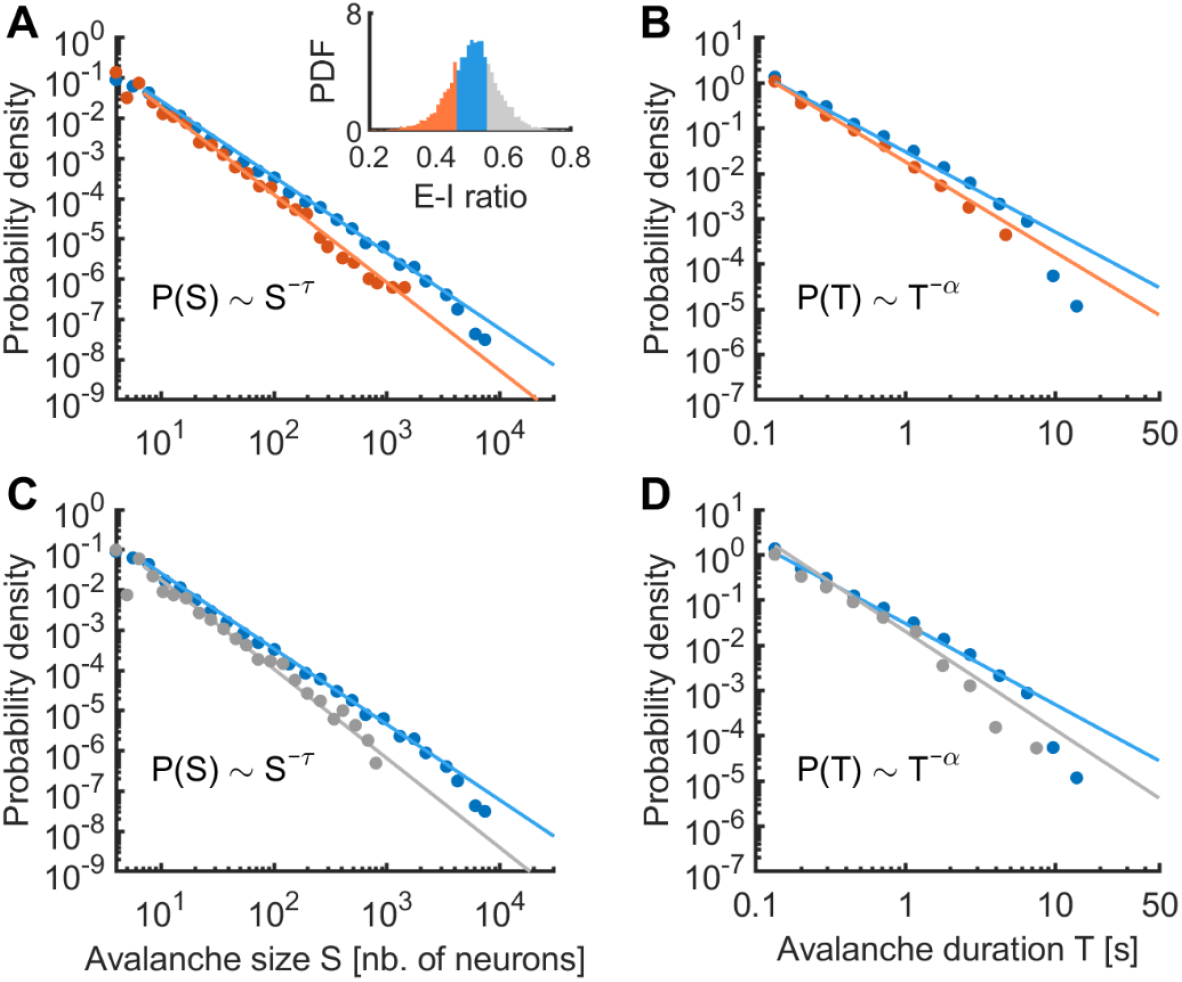
Model’s neuronal avalanche exponents as a function of the E-I ratio. The model was simulated at its critical point, for parameters Δ*w* = 0.1 and *w*_*I*_ = 7. The resulting distributions of avalanches sizes and duration were examined for different levels of E-I ratios. **A)** Distribution of avalanche sizes (*S*) for avalanches occurring at low E-I ratios (*red*, lowest 25% E-I ratios) and at balanced E-I ratios (*blue*, E-I ratios within the 25^th^-75^th^ percentiles). Solid lines indicate estimated power laws. *Inset:* distribution of E-I ratio. *Red:* low E-I ratios; *blue:* balanced E-I ratios; *gray*: high E-I ratios (highest 25% E-I ratios). **B)** Distribution of avalanche durations (*T*) for avalanches occurring at low (*red*) and balanced (*blue*) E-I ratios, respectively. Solid lines: estimated power laws. **C-D)** Same as (A) and (B) but comparing avalanches occurring at high E-I ratios (*gray*) and at balanced E-I ratios (*blue*).

### Supplemental tables

**Table S1.** Summary of the data and the statistics of the neuronal avalanches for each dataset. The table presents the number of neurons of each cell type (E, I, and Ch) and the statistics of neuronal avalanches. Avalanche size *S* and duration *T* distributions were fitted with truncated power laws (*P*(*S*)∼*S*^−*τ*^ and *P*(*T*)∼*T*^−*α*^) using MLE. Bootstrap resampling was employed to calculate the estimation error of the MLE power exponents^77^. Truncation cutoffs were [*S*_min_; *S*_max_] = [7; 10^4^] and [*T*_min_; *T*_max_] = [0.267 *s*; 30 *s*] for avalanche sizes and durations, respectively. The normalized log-likelihood ratio (LLR) compared power-law and log-normal fits, with significantly positive LLR values favoring the power-law model^78,79^; *: p < 0.01; **: p < 0.001. The relationship between the average avalanche size ⟨*S*⟩(*T*) and duration *T* was fitted with a power-law, ⟨*S*⟩(*T*)∼*T*^−1⁄*σνz*^, using least squares within the defined duration and size cutoffs to estimate the exponent *σνz* and its error. The power-law fits of ⟨*S*⟩(*T*) were compared the those obtained with an exponential function by calculating the ratio of the explained variances (REV) of the competing models. Ratios significantly greater than 1 support the power-law hypothesis (*: p < 0.01; **: p < 0.001).

| Fish<br># | Nb. of neurons | | | | $P(S) S^{-\tau}$ | | $P(T) T^{-\alpha}$ | | $\langle S \rangle(T) T^{1/\sigma \nu z}$ | |
| --- | --- | --- | --- | --- | --- | --- | --- | --- | --- | --- |
| | Total | E | I | Ch | $\tau$ | LLR | $\alpha$ | LLR | $\sigma \nu z$ | REV |
| 1 | 1943 | 1092 | 828 | 23 | $1.784 \pm 0.007$ | +12.5** | $2.711 \pm 0.025$ | +15.8** | $0.655 \pm 0.026$ | 1.10** |
| 2 | 2818 | 1542 | 1235 | 41 | $1.975 \pm 0.012$ | +14.6** | $2.708 \pm 0.037$ | +13.0** | $0.644 \pm 0.023$ | 1.15** |
| 3 | 1796 | 926 | 806 | 28 | $1.931 \pm 0.010$ | +16.4** | $2.643 \pm 0.031$ | +13.8** | $0.647 \pm 0.023$ | 1.21** |
| 4 | 2338 | 1308 | 1030 | 0 | $2.132 \pm 0.009$ | +10.6** | $3.310 \pm 0.042$ | +13.9** | $0.622 \pm 0.036$ | 1.10* |
| 5 | 2138 | 1201 | 909 | 28 | $2.173 \pm 0.015$ | +8.2** | $3.094 \pm 0.052$ | +11.1** | $0.667 \pm 0.018$ | 1.12** |
| 6 | 1756 | 1025 | 690 | 41 | $2.035 \pm 0.015$ | +11.3** | $2.783 \pm 0.047$ | +13.6** | $0.669 \pm 0.017$ | 1.11** |
| 7 | 2273 | 1240 | 1016 | 17 | $2.073 \pm 0.010$ | +18.9** | $2.880 \pm 0.030$ | +21.1** | $0.612 \pm 0.008$ | 1.16** |
| 8 | 2051 | 1092 | 945 | 14 | $1.977 \pm 0.014$ | +13.1** | $2.783 \pm 0.049$ | +10.5** | $0.721 \pm 0.074$ | 1.41** |
| 9 | 2622 | 1416 | 1179 | 27 | $1.956 \pm 0.010$ | +20.5** | $2.575 \pm 0.030$ | +15.0** | $0.601 \pm 0.015$ | 1.12** |
| 10 | 2265 | 1253 | 985 | 27 | $2.038 \pm 0.010$ | +15.0** | $2.761 \pm 0.033$ | +17.2** | $0.623 \pm 0.018$ | 1.11** |

**Table S2.** Summary of the data and the statistics of the neuronal avalanches using the spatially unconstrained avalanche definition. Neuronal avalanches were identified by summing the activity across all *N* recorded neurons at each time bin. An avalanche was defined as a period when this summed activity exceeded a threshold *θ* (0.5% of the neurons). The table indicates the exponents of the power laws (*P*(*S*)∼*S*^−*τ*^ and *P*(*T*)∼*T*^−*α*^) estimated using MLE^77^. No cutoffs were used. Asterisks indicate the significance of power-law fits using LLR comparisons of power-law and log-normal fits^78,79^; *: p < 0.01; **: p < 0.001. The average avalanche size ⟨*S*⟩(*T*) was fitted to a power law, ⟨*S*⟩(*T*)∼*T*^−1⁄*σνz*^, using least squares to estimate the exponent *σνz* and its error. The fit was compared to an exponential model by computing the ratio of explained variance (REV). Ratios >1 indicate support for the power law (*p < 0.01; **p < 0.001).

| Fish<br># | $P(S) S^{-\tau}$ | | $P(T) T^{-\alpha}$ | | $\langle S \rangle(T) T^{1/\sigma\nu z}$ | |
| --- | --- | --- | --- | --- | --- | --- |
| | $\tau$ | LLR | $\alpha$ | LLR | $\sigma\nu z$ | REV |
| 1 | $1.335 \pm 0.008$ | -10.8<sup>n.s.</sup> | $1.594 \pm 0.013$ | +1.3<sup>n.s.</sup> | $1.094 \pm 0.023$ | 1.28<sup>**</sup> |
| 2 | $1.694 \pm 0.014$ | +11.3<sup>**</sup> | $1.730 \pm 0.014$ | +7.1<sup>**</sup> | $0.862 \pm 0.014$ | 1.29<sup>**</sup> |
| 3 | $1.556 \pm 0.013$ | +9.2<sup>**</sup> | $1.637 \pm 0.014$ | +6.9<sup>**</sup> | $0.859 \pm 0.021$ | 1.34<sup>**</sup> |
| 4 | $1.517 \pm 0.015$ | -3.2<sup>n.s.</sup> | $1.587 \pm 0.011$ | -1.4<sup>n.s.</sup> | $0.931 \pm 0.012$ | 1.31<sup>*</sup> |
| 5 | $1.839 \pm 0.013$ | +11.4<sup>**</sup> | $1.774 \pm 0.012$ | +2.6<sup>*</sup> | $0.847 \pm 0.010$ | 1.31<sup>**</sup> |
| 6 | $1.687 \pm 0.013$ | +9.5<sup>**</sup> | $1.675 \pm 0.014$ | +2.9<sup>*</sup> | $0.865 \pm 0.012$ | 1.27<sup>**</sup> |
| 7 | $1.560 \pm 0.013$ | +8.8<sup>**</sup> | $1.568 \pm 0.013$ | +2.6<sup>*</sup> | $0.831 \pm 0.013$ | 1.29<sup>**</sup> |
| 8 | $1.669 \pm 0.013$ | +15.4<sup>**</sup> | $1.650 \pm 0.014$ | +5.2<sup>**</sup> | $0.786 \pm 0.011$ | 1.29<sup>**</sup> |
| 9 | $1.588 \pm 0.012$ | +13.2<sup>**</sup> | $1.558 \pm 0.013$ | +4.1<sup>**</sup> | $0.831 \pm 0.013$ | 1.32<sup>**</sup> |
| 10 | $1.700 \pm 0.013$ | +16.5<sup>**</sup> | $1.652 \pm 0.012$ | +6.2<sup>**</sup> | $0.814 \pm 0.016$ | 1.26<sup>**</sup> |

### Supplemental notes

#### Note S1: Analysis of pairwise correlations as a function of cell type

We examined the stationary correlations of collective activity as a function of cell types. We calculated the pairwise correlations between different pairs of cell types (**Supplemental Figure S4A**). The average correlation between neurons was low (0.02) and was not significantly different for different pairs of cell types (one-way ANOVA: *F*_(14,42)_ = 1.42, *p* = 0.185). Furthermore, we investigated whether different cell types tend to form clusters of correlated neurons by analyzing the modular structure of the correlation matrix as a function of cell types using a community detection algorithm (see Supplemental Methods). A mild modular structure was observed (**Supplemental Figure S4B**), with low modularity indices ranging between 0.158 and 0.346 across the different fish (**Supplemental Figure S4C**). These modules, however, were not predominantly composed of a single cell type. To quantify this, we calculated the ratio *χ* between the proportion of neurons of type *i* (with *i* = E, I, or Ch) in the detected modules and the proportion expected in random partitions of the correlation matrix of equal size. This ratio remained close to 1 for all cell types (its average ranged between 0.96 and 1.09 for the different cell types), indicating that the modules did not show a preference for specific cell types (**Supplemental Figure S4D**). Lastly, we analyzed the topological positions of neurons of different cell types within the network defined by the correlation matrix. Specifically, we calculated the betweenness centrality of neurons, a measure of node centrality in a graph or network (see Supplemental Methods). The average betweenness centrality did not differ significantly between cell types (one-way ANOVA: *F*_(2,26)_ = 2.52, *p* = 0.10; **Supplemental Figure S4E**). Overall, these results indicate that the structure of stationary correlations was not straightforwardly organized as a function of cell types.

#### Note S2: Shape collapse of neuronal avalanches

Close to criticality, the average avalanche profile, ⟨*S*(*t, T*)⟩, is expected to exhibit scale invariance across temporal scales^5,13,43^. Specifically, when time is normalized by the avalanche duration, the scaled avalanche profile follows a universal form: ⟨*S*(*t, T*)⟩*T*^−*a*^ = *F*(*t*⁄*T*), that is independent of the avalanche duration *T*. This scale invariance is commonly referred to as “shape collapse”. To estimate the scaling parameter *a*, we used the method of Marshall et al.^77^ which determines the value of *a* that yields the optimal collapse of avalanche profiles across durations. Specifically, the method estimates the scaling parameter *a* that minimizes the variance 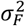 across the avalanche profiles in the normalized time (*t*/*T*). Near criticality, the scaling parameter *a* is expected to be related to the exponent *σνz* via: *a* = *σνz*^−1^ − 1. This is the consequence of the relationship ⟨*S*⟩(*T*)∼*T*^1⁄ *σνz*^ and ⟨*S*⟩(*T*) being equal to the integral of ⟨*S*(*t, T*)⟩ = *T*^*a*^*F*(*t*⁄*T*), over the interval *t* ∈ [0, *T*]. Shape collapse therefore provides an independent and complementary approach to both assess criticality and estimate the exponent *σνz*. The empirical avalanche profiles exhibited a stereotyped buildup and termination over a wide range of durations (**Supplemental Figure S5A–B**), such that the average time course of short avalanches closely resembled that of longer ones. This behavior was observed for avalanche profiles computed from the joint excitatory and inhibitory activity, ⟨*S*(*t, T*)⟩ (**Supplemental Figure S5A-B**, *top*), as well as when excitatory and inhibitory components were analyzed separately, ⟨*S*_*E*_(*t, T*)⟩ and ⟨*S*_*I*_(*t, T*)⟩ (**Supplemental Figure S5A-B**, *middle* and *bottom*). Notably, the scaling exponents estimated from profile collapse were consistent with the values of *σνz* obtained independently from the scaling relation between ⟨*S*⟩ and *T*, for the different types of avalanche profiles (**Supplemental Figure S5C**).

## Supplemental methods

### Graph theory measures

We used a community detection algorithm to identify modules in the correlation matrix *C* of the neuronal activity, representing functional connectivity. The algorithm, which handles weighted, signed matrices, is described in Rubinov and Sporns (2010) [S1] and its implementation is available in the Brain Connectivity Toolbox (https://sites.google.com/site/bctnet/). Briefly, the method employs the Louvain community detection algorithm [S2] to find the optimal community structure, which divides the network into non-overlapping groups of nodes, maximizing within-group connections and minimizing between-group connections. The modularity index *Q* measures the degree of modular segregation in the graph. *Q* values close to 1 indicate low overlapping modules, while *Q* values close to 0 indicate that inter-module connections are comparable to those in a random graph. *Q* values larger than 0.3–0.4 are usually interpreted as evidence that the subgraphs of the corresponding partition constitute distinct modules [S3].

We also measured the centrality of neurons within the network given by the correlation matrix of the neuronal activity (*C*). For this, we measure the betweenness centrality as implemented in the Brain Connectivity Toolbox. Briefly, node betweenness centrality is the fraction of shortest paths in the network that contain a given node. To calculate the betweenness centrality, we first binarized the matrix *C* by detecting those correlations that were significantly different from zero. This was done by comparing *C* to the correlation matrix of randomized data, noted *C*^sh^, obtained after shuffling in time the series of neuronal activations independently for each neuron (100 repetitions). We next defined the binary graph *G* as follows: *G*_*ij*_ = 1 if the functional correlation between neurons *i* and *j, C*_*ij*_, was larger than three standard deviations of the corresponding entries of the correlation matrix of randomized data 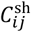, i.e., 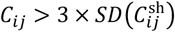, and *G*_*ij*_ = 0 otherwise. Finally, the normalized betweenness centrality of each node of graph *G* was given as:

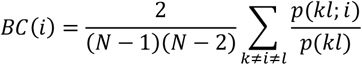

where *p*(*kl*) is the total number of shortest paths in graph *G* from node *k* to node *l, p*(*kl*; *i*) is the number of those paths that pass through the node *i*, and *N* is the number of neurons. The betweenness centrality is normalized by dividing by the number of pairs of nodes of an undirected graph of *N* nodes, i.e., 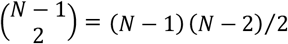.

